# Neonatal microbiota colonization drives maturation of primary and secondary goblet cell mediated protection in the pre-weaning colon

**DOI:** 10.1101/2024.07.03.601781

**Authors:** Åsa Johansson, Mahadevan Venkita Subramani, Bahtiyar Yilmaz, Elisabeth Nyström, Elena Layunta, Liisa Arike, Felix Sommer, Philip Rosenstiel, Lars Vereecke, Louise Mannerås Holm, Andy Wullaert, Thaher Pelaseyed, Malin E.V. Johansson, George M.H. Birchenough

**Affiliations:** Department of Medical Biochemistry & Cell Biology, Institute of Biomedicine, University of Gothenburg, Sweden; Wallenberg Centre for Molecular & Translational Medicine, University of Gothenburg, Sweden; Department for BioMedical Research, University of Bern, Switzerland; Institute of Clinical & Molecular Biology, University of Kiel, Germany; VIB-UGent Center for Inflammation Research, Ghent, Belgium; Department of Internal Medicine and Pediatrics, Ghent University, Ghent, Belgium; Wallenberg Laboratory, Institute of Medicine, University of Gothenburg, Sweden; Department of Biomedical Sciences, University of Antwerp, Belgium

## Abstract

In the distal colon, mucus secreting goblet cells primarily confer protection from luminal microorganisms via generation of a sterile inner mucus layer barrier structure. Bacteria-sensing sentinel goblet cells provide a secondary defensive mechanism that orchestrates mucus secretion in response to microbes that breach the mucus barrier. Previous reports have identified mucus barrier deficiencies in adult germ-free mice, thus implicating a fundamental role for the microbiota in programming mucus barrier generation. In this study, we have investigated the natural neonatal development of the mucus barrier and sentinel goblet cell-dependent secretory responses upon postnatal colonization. Combined *in vivo* and *ex vivo* analyses of pre- and post-weaning colonic mucus barrier and sentinel goblet cell maturation demonstrated a sequential microbiota-dependent development of these primary and secondary goblet cell-intrinsic protective functions, with dynamic changes in mucus processing dependent on innate immune signalling *via* MyD88, and development of functional sentinel goblet cells dependent on the NADPH/Dual oxidase family member Duox2. Our findings therefore identify new mechanisms of microbiota-goblet cell regulatory interaction and highlight the critical importance of the pre-weaning period for the normal development of colonic barrier function.

## Introduction

The intestinal mucosal epithelium is protected from luminal microorganisms by dynamic mucus barrier structures secreted by epithelial goblet cells (GCs) (*1*). The polymeric gel-forming Mucin-2 (Muc2) glycoprotein is the core structural component of mucus, and its genomic deletion results in increased susceptibility to infection (*2*), inflammation (*3*) and tumorigenesis (*4*), underlining the crucial role of mucus as a legislator of intestinal homeostasis.

The greatest environmental challenge faced in the intestine is containment of the colonic microbiota, a dense microbial community that can incorporate many opportunistic pathogens. The colonic mucus system has evolved to deal with this challenge by forming a dense, tissue-adherent inner mucus layer (IML) barrier that is impenetrable to most microbes (*5*). The Muc2 polymers that form the IML are protected from microbial degradation by heavy *O*-glycosylation (*6*), intermolecular mucin isopeptide crosslinking (*7*) and the presence of proteins that aggregate or kill mucus invasive bacteria (*8–10*). IML Muc2 polymers undergo post-secretory processing by endogenous proteases (*11*) that expand the polymeric network, and likely contributes towards faecal encapsulation of the colonic microbiota (*12*). Consequently, the IML is the primary GC-intrinsic protective system in the colon and its protective functions are strongly influenced by several layers of post-translational Muc2 modification.

Besides IML formation, colonic GCs have additional proposed roles in mucosal protection. A bacteria-sensing subpopulation of upper crypt GCs referred to as sentinel GCs (senGCs) orchestrates defensive mucus secretion in response to elevated levels of specific Toll-like receptor (TLR) ligands *via* assembly of an NLR family pyrin domain containing 6 (Nlrp6)-dependent inflammasome complex (*13*). This process is believed to function as a secondary protective function that guards colonic crypts against bacteria that have penetrated the IML. Furthermore, GCs can act as GC-associated antigen passages (GAPs) to transfer luminal material to lamina propria immune cells and thereby regulate mucosal immunity against dietary and microbial antigens (*14, 15*), thus illustrating that GCs have protective functions beyond constitutive mucus secretion.

While knowledge of GC protective functions continues to expand, our understanding of how these systems are established during early life development remains incomplete. Postnatal mucus barrier development in the small intestine varies along the proximal-distal axis, but the regulatory mechanisms governing this heterogeneity are unknown (*16*). Prior work focused on the large intestine indicates that colonic GAPs are induced during the pre-weaning phase and repressed during the suckling-weaning transition (*17*); however, postnatal IML or senGC functional maturation has not been described. Studies in adult germ-free (GF) mice indicate that IML functions are compromised in the absence of a microbiota (*12, 18, 19*), thus hinting that these protective functions are likely to develop during establishment of the colonic microbiota in the neonatal-weaning period. We therefore sought to define the postnatal developmental dynamics of colonic IML and senGC maturation and examine their relation to microbiota colonization in order to provide a more complete understanding of defensive maturation in the large intestine.

## Results

### The colonic inner mucus layer matures in the pre-weaning environment

The colonic IML is the primary GC-intrinsic protective system (*5*). The adult IML is a tissue adherent structure that is impenetrable to bacteria-sized objects and undergoes post-secretion processing driven in part by the action of endogenous metalloproteases (*11*). Characterization of postnatal IML development over the neonatal, weaned and adult developmental stages was determined by *ex vivo* analysis of live colonic tissues obtained from Wistar rats ranging in age from 1 day *postpartum* (P1)-P15 (neonatal stage), P22-P30 (weaned stage) and >P100 (adult stage, 14 week or older animals) (**Fig. 1A**). Rats were used in this case to provide sufficiently large tissue for robust *ex vivo* IML analysis at younger ages that would not be possible with mouse tissues. A tissue adherent mucus layer was observed at all ages; however, we noted a gradual increase in IML thickness from P1 (∼50 µm) to P15 (∼100 µm) that was subsequently maintained into adulthood (**Fig. 1B**). Strikingly, analysis of IML barrier function did not follow a gradual pattern, with a sharp transition from a penetrable IML structure in very young (P1-2) pups to an impenetrable, adult-like IML at P3 (**Fig. 1C, D**). We confirmed postnatal IML formation *in vivo* by immunostaining tissue sections for the critical IML structural protein Muc2. This analysis identified a well-preserved, stratified IML structure encapsulating the faecal microbiota at P3 and older colonic tissues that was not found at P2 (**Fig. 1E**). Notably, we did not observe any direct evidence of microbiota-epithelial contact at P2, despite the lack of a well-developed IML at this age. Lastly, we assessed post-secretion processing of the IML by quantification of the *ex vivo* mucus growth rate. This analysis identified variable alterations during the neonatal period, with a rate of ∼1 µm/min observed immediately *postpartum* (P1), reducing to ∼0.5 µm/min from P2-P4 before gradually increasing to a rate of ∼2 µm/min between P5-P10 that was subsequently maintained into adulthood (**Fig. 1F**). Combined, these analyses indicate that IML formation principally occurs in the pre-weaning period, with dynamic and heterogenous alterations in IML thickness, barrier function and processing in the early postnatal environment.

**Figure 1:**
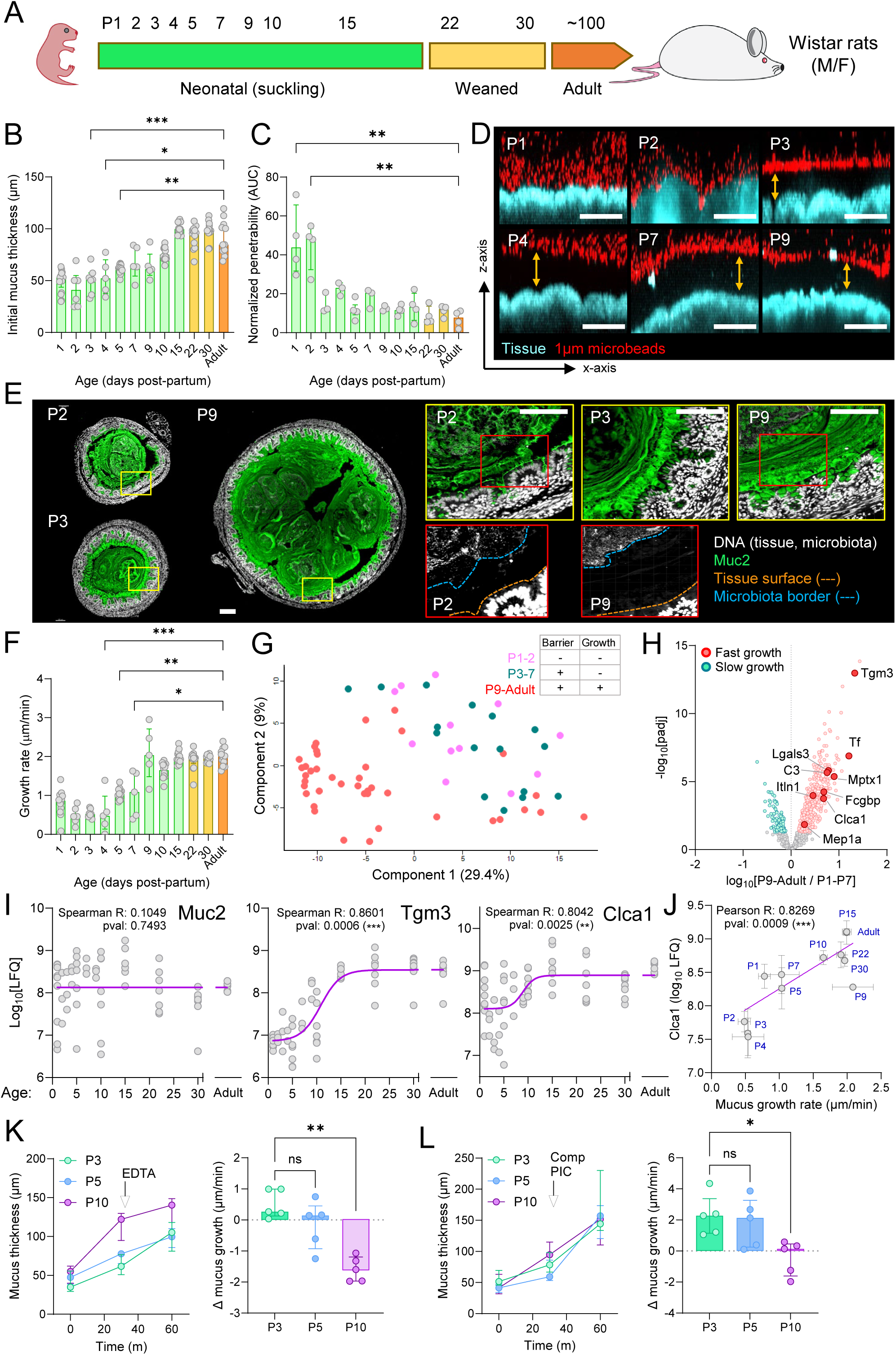
Postnatal maturation of the colonic inner mucus layer barrier. **A)** Sampling timepoints in days *post-partum* (P) of neonatal, weaned and adult Wistar rats. **B)** IML thickness quantified by *ex vivo* needle measurement. **C)** IML barrier function quantified by *ex vivo* microbead penetration. **D)** Representative confocal z-stacks used to generate data shown in (C). Images show x/z-axis cross-sections of colonic tissues (blue) overlaid with microbeads (red). Impenetrable mucus indicated (yellow arrows). **E)** Confocal micrographs of fixed colonic tissue sections stained for DNA (grey) and Muc2 (green). Dashed lines in high magnification P2 and P9 images show tissue surface (orange) and microbiota border (blue). **F)** IML growth rate quantified by *ex vivo* needle measurement. **G)** Principle component analysis plot of complete mucus proteome data. Samples colour coded based on age-dependent IML *ex vivo* phenotype: P1-P2 (low barrier function, slow growth; pink), P3-P7 (high barrier function, slow growth; teal), P9-Adult (high barrier function, fast growth; red). **H)** Volcano plot comparing complete mucus proteome data from slow growth (P1-P7) and fast growth (P9-Adult) samples. Proteins significantly enriched in slow (teal) or fast (red) growth groups are indicated. **I)** Abundance of Muc2 (left), Tgm3 (middle) and Clca1 (right) proteins at different ages from label free quantification (LFQ) of complete mucus proteome data. Non-linear regression curves (purple) are indicated for each dataset. Spearman correlation R and p values between age and protein LFQ are shown. **J)** Correlation of *ex vivo* mucus growth rate and Clca1 mucus proteome abundance (LFQ) at different ages. Simple linear regression (purple) is indicated. Pearson correlation R and p value between mucus growth rate and Clca1 LFQ is shown. **K)** *Ex vivo* mucus growth and response to metalloprotease inhibition by EDTA treatment at different ages. Graphs show mucus thickness values (left) and changes in mucus growth rate (Δ mucus growth; right) in response to EDTA. **L)** *Ex vivo* mucus growth and response to serine/cysteine protease inhibition by Complete Protease Inhibitor Cocktail (Comp PIC) treatment at different ages. Graphs show mucus thickness values (left) and changes in mucus growth rate (Δ mucus growth; right) in response to Comp PIC. Data represents n=5-19 (A-J) or n=5 (K-L) animals per group, as indicated. All data is pooled from at least 3 independent litters. All histograms show median and interquartile range. Statistical comparisons between groups by Kruskal Wallis and Dunn’s multiple comparison (B, C, F, K, L) or Welch’s t-test and Benjamini-Hochberg FDR correction (H); p<0.05 (*), <0.01 (**) <0.001 (***), <0.0001 (****). Image scale bars are 100µm.

### Postnatal IML maturation reflects mucus compositional and processing dynamics

We subsequently sought to identify potential molecular drivers of postnatal IML maturation. Compositional analysis of the complete IML proteome by label-free mass spectrometry indicated significant variability in mucus protein composition at different ages. Principal component analysis (PCA) of proteomic data and grouping of samples into three phenotypic groups covering P1-2 (penetrable, low growth), P3-P7 (impenetrable, low growth) and P9-adult (impenetrable, high growth) age ranges identified sample clustering associated with mucus growth rate (but not barrier function) phenotype (**Fig. 1G**). Pooled comparison of fast growth (P9-adult) and low growth (P1-7) samples detected significant enrichment of multiple proteins in the high growth group, including proteins with established or likely roles in antimicrobial defence (e.g. Itln1, Mptx1, Lgals3, Tf, C3) (**Fig. 1H**). The abundance of Muc2 itself was not significantly altered during postnatal development; however, the abundance of two key Muc2 modifying enzymes with known roles in promoting IML integrity (Transglutaminase 3 – Tgm3) (*7*) and proteolytic processing (Calcium-activated chloride channel regulator 1 – Clca1) (*11*) both showed significant positive, monotonic correlations with age (**Fig. 1I**).

Clca1 is a critical regulator of the proteolytic IML expansion that drives *ex vivo* mucus growth (*11*), and Clca1 mucus abundance displayed positive linear correlation to mucus growth rates across different age groups (**Fig. 1J**). We therefore sought to clarify the role of Clca1 in postnatal IML processing dynamics by treating P3, P5 and P10 tissues with EDTA, which has been previously shown to inhibit mucus growth in a Clca1-dependent manner (*11*). While EDTA treatment had no impact on low growth samples (P3/5), a clear inhibitory effect was observed in high growth (P10) samples, thus indicating that the postnatal increase in growth was dependent on increased mucus processing by Clca1 (**Fig. 1K**). In order to validate the specificity of this approach, we also treated tissues with the pan-serine/cysteine inhibitor cocktail Complete (Comp PIC). As previously reported in adult mice (*11*), Comp PIC treatment had no impact on Clca1-dependent P10 mucus growth; however, inhibitor treatment resulted in an unexpected increase in mucus growth in P3/5 tissues, indicating that mucus growth is actively suppressed by protease function in these samples (**Fig. 1L**).

Together, these experiments demonstrated that dynamic alterations in postnatal IML properties are likely linked to the increased abundance of specific Muc2 processing proteins (e.g. Clca1) and that this coincides with increased levels of proteins that reinforce mucus barrier integrity (Tgm3) and antimicrobial functions. Surprisingly, our data also indicated that the low mucus growth observed in the early neonatal period is a result of serine/cysteine protease activity, indicating the unexpected presence of active mucus growth suppression in early neonatal tissues.

### Microbiota colonization and MyD88 signalling regulate specific aspects of IML maturation

Previous investigations comparing germ-free (GF) and conventionally raised (ConvR) mice have established that IML function and composition can be regulated by the intestinal microbiota (*18–20*); however, as these comparisons have exclusively used adult mice it remains unclear if they reflect the mechanistic basis of natural, postnatal IML formation. To address this question, we therefore compared neonatal colonic tissue samples from ConvR and GF C57BL/6 mice over the P1-P9 period where we had observed IML maturation in rats using an *ex vivo* approach (**see Fig. 1**). In order to confirm that both neonatal mice and rats are exposed to similar post-natal colonization dynamics, we initially quantified microbiota colonization using 16S qPCR to detect both total and taxon-specific (Proteobacteria, Firmicutes, Bacteroidetes) bacterial load in stool samples collected from prenatal and postnatal colonic tissues. In both rats and mice, bacterial colonization was not clearly observed until P1-P2, with initial colonization driven by high abundance of the Firmicutes and Proteobacteria phyla, compared to the Bacteroidetes-dominated adult samples (**Fig. 2A**). This delay between birth and microbiota establishment was confirmed by 16S FISH staining of luminal content from neonatal mice (**Fig. 2B**), indicating the establishment of the IML barrier observed in rats between P2-P3 occurred immediately after initial microbiota seeding of the colonic environment.

**Figure 2:**
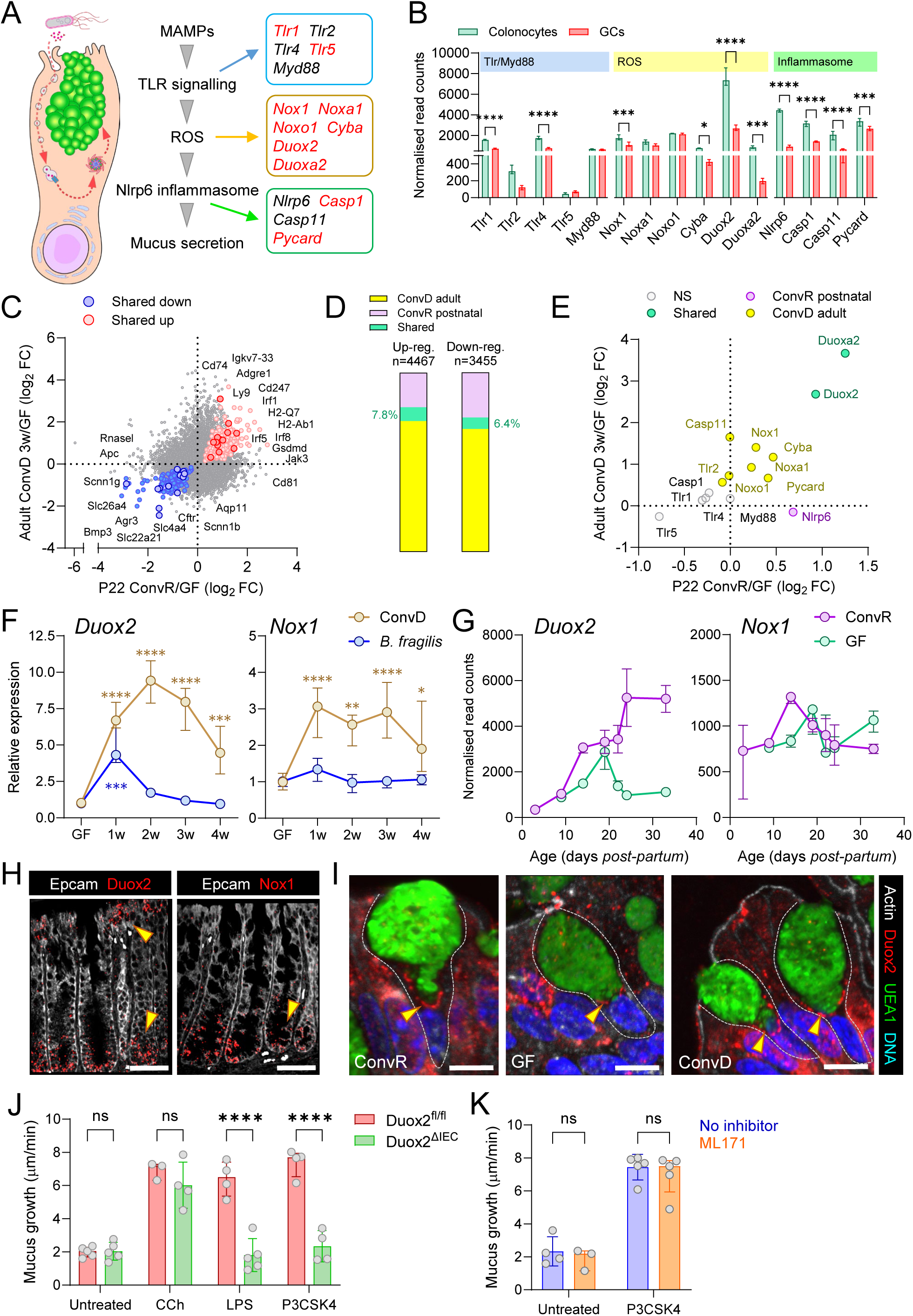
Postnatal IML maturation is driven by both microbiota-dependent and independent factors. **A)** Total and taxon-specific colonic bacterial load in foetal and postnatal Wistar rats (left) and C57BL/6 mice (right) quantified by 16S qPCR of stool DNA. IML formation period between P2-P3 in rats is indicated. **B)** Confocal micrographs of fixed mouse colonic tissue stained for luminal bacteria by 16S FISH. **C)** Confocal micrographs of fixed colonic tissue sections stained for DNA (grey) and Muc2 (green). Dashed lines in high magnification images show tissue surface (orange) and IML border (yellow). Stratified Muc2 layers present in ConvR but not GF P3 IML are indicated (arrows). **D)** Comparison of gene expression ratios between ConvR P14:P3 mice (age-dependent) and P14 ConvR:GF mice (microbiota-dependent) quantified by DESeq2 analysis of bulk colonic RNA sequencing data. Genes significantly regulated by age (orange) or by both age and colonization status (blue) are indicated. **E)** *Ex vivo* mucus growth in GF, ConvD or ConvR tissue before (pre) and after (post) serine/cysteine protease inhibition by Complete Protease Inhibitor Cocktail (Comp PIC) or the cysteine protease-specific inhibitor E64. **F)** *Ex vivo* analysis of *MyD88*^+/+^ and *MyD88*^-/-^ IML thickness and barrier function by microbead penetration. Images are x/z-axis cross-sections of confocal z-stacks of showing colonic tissues (grey) overlaid with microbeads (red). Impenetrable mucus indicated (yellow arrows). **G)** *Ex vivo* analysis of *MyD88*^+/+^ and *MyD88*^-/-^ IML thickness based on images shown in (F). **H)** *Ex vivo* analysis of *MyD88*^+/+^ and *MyD88*^-/-^ IML barrier function based on images shown in (F). **I)** *Ex vivo* mucus growth in *MyD88*^+/+^ and *MyD88*^-/-^ tissue in response to metalloprotease (EDTA) or serine/cysteine protease (Comp PIC) inhibition. Data represents n=4-5 (A-C, E, I), n=6-8 (F-H) or n=3 (D) animals per group, as indicated. All data is pooled from at least 2 independent litters or experiments. All histograms show median and interquartile range. Statistical comparisons between groups by DESeq2 (D), 2-way ANOVA and Fishers LSD (E, I) or Mann Whitney test (G, H); p<0.05 (*), <0.01 (**) <0.001 (***), <0.0001 (****). Image scale bars are 50µm (B) or 100µm (C).

As *ex vivo* analytical approaches are not suited to neonatal mice due to tissue size limitations, IML formation was instead examined histologically *via* immunostaining for Muc2 in tissue sections obtained from both ConvR and GF neonatal mice. Comparable levels of luminal Muc2 were observed in both groups at all ages, and Muc2 staining at P2 did not show any clear organisation (**Fig. 2C**); however, at P3 and older a stratified IML barrier structure encapsulating the microbiota was observed in ConvR mice, which was absent in age-matched GF mice. Consequently, our data indicated that the rapid postnatal development of the colonic IML is dependent on microbiota colonization, and that developmental kinetics observed by *ex vivo* analysis of rats (**Fig. 1C, D**) is similar to that observed in mice.

We have previously established that EDTA inhibits GF mucus growth to the same extent observed in ConvR tissues, indicating that Clca1-dependent mucus processing is independent of microbiota colonization (*11*). In order to determine if this was consistent during postnatal development, we performed bulk sequencing of mRNA extracted from ConvR P3 and P14 tissues and GF P14 tissues. We then compared whole transcriptome expression ratios between ConvR P14:P3 and P14 ConvR:GF samples to identify genes that showed either age and/or colonization dependent changes in expression (**Fig. 2D**). Notably, genes encoding proteins with age-dependent increases in mucus abundance linked to Muc2 processing (Clca1, Tgm3) and antimicrobial activity (Itln1, Mptx1, Lgals3, Tf) were significantly induced by age but not affected by colonization status, indicating that these expression changes related to IML maturation were developmentally hard-wired, independent of microbiota colonization.

We next sought to determine whether the serine/cysteine protease suppression of mucus growth observed in early neonates (**Fig. 1L**) was a microbiota-regulated feature of IML maturation. Similar to the early neonatal phenotype, *ex vivo* analysis of adult GF colonic tissue demonstrated increased mucus growth in response to both serine/cysteine protease inhibition (Comp PIC) and specific inhibition of cysteine protease activity using the inhibitor E64 (**Fig. 2E**). Accordingly, full microbiota conventionalisation (ConvD) of GF mice for 4 weeks restored insensitivity to E64 treatment as observed in ConvR mice (**Fig. 2E**), thus indicating that proteolytic suppression of mucus processing is a microbiota-regulated phenomenon.

Prior studies have implicated microbiota recognition *via* the TLR-MyD88 signalling axis as an important regulator of colonic barrier function, including expression of Muc2 (*21, 22*). We therefore investigated the possibility that MyD88 function was critical for normal IML maturation by *ex vivo* comparison of mucus phenotypes from adult *MyD88*^+/+^ and *MyD88*^-/-^ mice. No significant differences between wild-type and knockout littermate mice were observed in terms of baseline IML thickness or barrier function, indicating that IML formation is independent of MyD88 function (**Fig. 2F-H**). In terms of mucus processing, baseline growth rate or sensitivity to EDTA-mediated metalloprotease inhibition were unaffected by MyD88 deficiency; however, treatment of *MyD88*^-/-^ tissue with Comp PIC induced increased mucus growth rates comparable to those observed in early neonatal tissues (**Fig. 2I**). This demonstrated that MyD88 signalling does not regulate Clca1-mediated mucus expansion, but is a critical negative regulator of cysteine protease-mediated mucus growth suppression.

Combined, these experiments demonstrate that natural, postnatal maturation of IML barrier properties and mucus processing are regulated by a combination of both microbiota-dependent and independent mechanisms, and that MyD88 innate immune signalling plays an important role in dampening proteolytic suppression of mucus expansion in early life.

### Microbiota colonization drives postnatal goblet cell maturation

Having examined the role of the microbiota in postnatal IML maturation, we next sought to establish how both age and microbial exposure influenced the postnatal maturation dynamics of the mucus secreting colonic goblet cell (GC) population. Accordingly, gene expression data from bulk sequencing of RNA extracted from neonatal (P9-P22), weaned (P24-P33) and adult (>P100) ConvR and GF colonic tissues (**Fig. 3A**) was filtered for 4098 genes previously shown to have significantly enriched expression in colonic GCs (*23*). Comparison of adult ConvR and GF expression data using DESeq2 identified microbial modulation of 120 GC-enriched genes, including induction of the important ER chaperone BIP (*Hspa5*) and repression of the secretory cell-regulating enzyme Ido1 (**Fig. S1A**). While these differences were potentially relevant to GC function, only 12/120 genes had a log_2_ fold-change ≥1, reflecting only modest microbiota-dependent expression differences in adult tissues.

**Figure 3:**
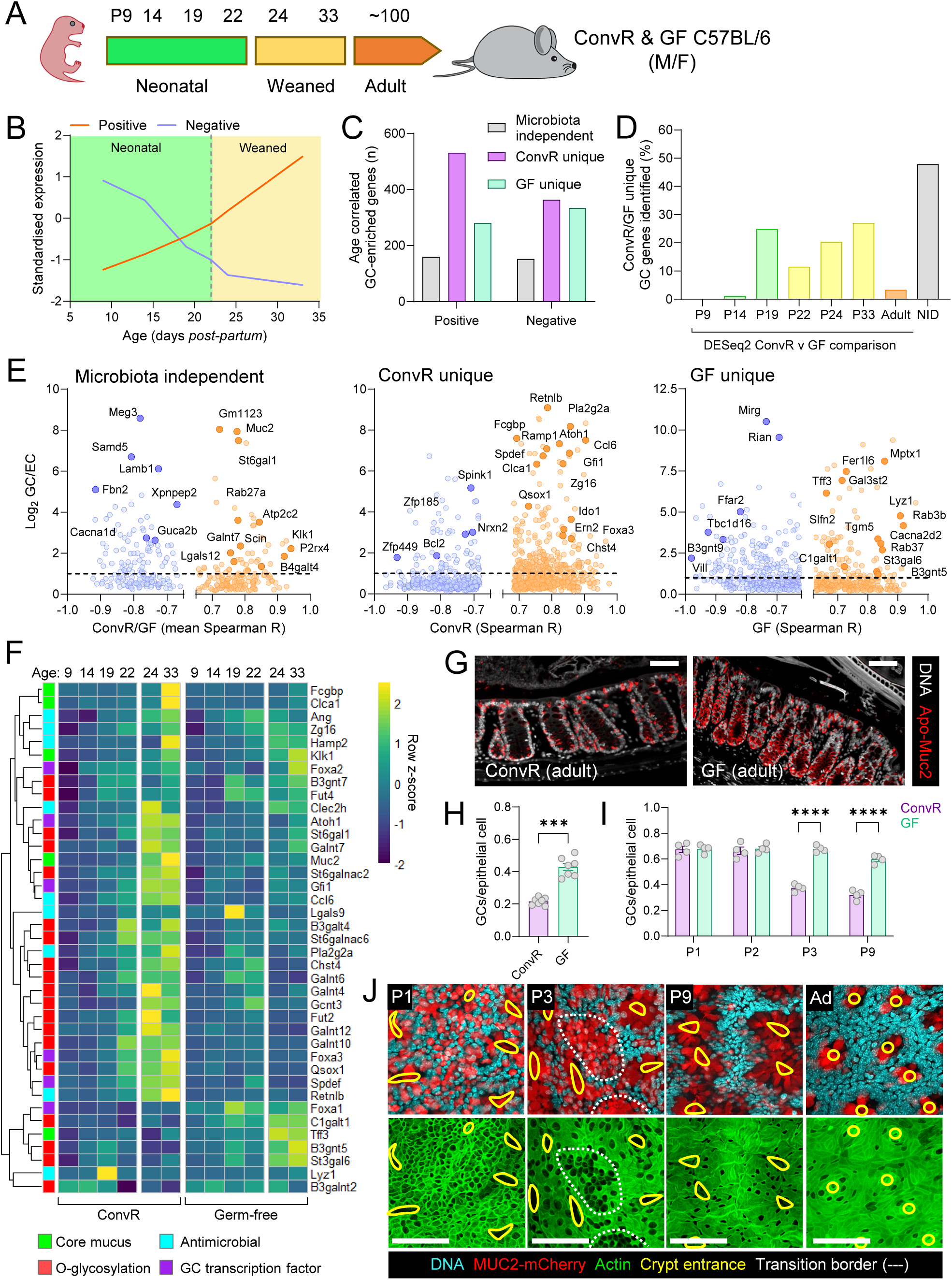
Microbiota colonization and goblet cell maturation. **A)** Sampling timepoints in days *post-partum* (P) of neonatal, weaned and adult ConvR and GF mice. **B)** Illustration of postnatal gene expression patterns with positive or negative monotonic correlation to age. **C)** Total number of GC-enriched genes with positive or negative monotonic correlation to age in both ConvR and GF mice (microbiota-independent) or in only ConvR or GF mice (microbiota-dependent) determined by Spearman correlation analysis of DESeq2 normalized RNA sequencing data. **D)** Proportion of microbiota-dependent gene expression patterns shown in (C) identified by standalone DESeq2 comparison of ConvR and GF samples at individual ages. Genes not identified in any pairwise comparison classified as “not identified” (NID). **E)** Comparison of Spearman correlation coefficients (R) and GC expression enrichment (log_2_ goblet cell:enterocyte expression ratios) of genes with significant positive (orange) or negative (blue) monotonic correlation to age. Plots show GC-enriched genes with microbiota-independent (left) or -dependent (middle, right) expression patterns. **F)** Heatmap showing standardized expression (z-score) of selected GC-enriched genes with significant microbiota-dependent or independent expression correlation to age. **G)** Confocal micrographs of fixed colonic tissue from adult ConvR and GF mice stained for DNA (grey) and Apo-Muc2 (red). **H)** Quantification of Apo-Muc2 positive cells as a fraction of total epithelial cells in adult ConvR and GF tissue based on images shown in (G). **I)** Quantification Apo-Muc2 positive cells as a fraction of total epithelial cells in P1-9 neonatal ConvR and GF tissues based on images shown in Fig. S1C. **J)** Whole mount confocal imaging of neonatal and adult RedMUC2^98tr^ colon stained for DNA (blue), F-actin (green) and MUC2-mCherry (red). Crypt entrances (yellow line) and intercrypt transition zone between high and low GC density areas (white dashed line) are indicated. Images show x/y-axis maximum intensity projections. Data represents n=2-4 (C-F), n=4-7 (G-I) or n=4 (J) animals per group, as indicated. All data is pooled from at least 2 independent litters or experiments. All histograms show median and interquartile range. Statistical comparisons between groups by Spearman correlation with Benjamini-Hochberg FDR correction (C-F), Mann Whitney test (H) or 2-way ANOVA and Fishers LSD (I); p<0.05 (*), <0.01 (**) <0.001 (***), <0.0001 (****). Image scale bars are 50µm.

We hypothesised that the relatively minor differences in GC enriched gene expression between adult ConvR and GF mice might be related to the fact that GF mice are still exposed to non-viable microbial products in their food, which might eventually mask microbiota-dependent gene regulation differences occurring in the postnatal period. We further hypothesised that analysis of age-correlated gene expression patterns could detect differences that may not be apparent by pairwise comparison of ConvR and GF tissue at individual postnatal time-points. We therefore quantified gene expression in P9-P33 (neonatal-weaned) RNA samples and correlated normalised gene counts to days *postpartum* by calculation of Spearman’s rank coefficient (R) to identify significant positive or negative monotonic relationships between gene expression and age (**Fig. 3B**). Using this approach, we identified 1819 GC genes with age correlated gene expression, of which 83% were dependent on microbiota-colonization status (i.e. were uniquely detected in either ConvR or GF datasets) (**Fig. 3C**). Notably, comparison of these genes to those detected by DESeq2 analysis of ConvR and GF data at each age demonstrated that 48% were not detected at any age by individual pairwise comparison, thus illustrating the potential benefit of this approach (**Fig. 3D**). This indicated that microbial colonization exerts a significant regulatory effect on postnatal GC gene expression that is not captured by comparative analysis of adult ConvR and GF tissues.

We next examined the GC-enriched genes identified by our analysis. Comparison of individual gene Spearman R values and relative enrichment in GCs compared to enterocytes (ECs) illustrated that a number of highly enriched GC genes displayed age-correlated expression patterns (**Fig. 3E**). Notably, expression of the core mucus component *Muc2* showed a positive correlation with postnatal age that occurred in both ConvR and GF mice, whereas age-dependent induction of other highly abundant core components was uniquely observed in either ConvR (*Clca1*, *Fcgbp*, *Zg16*) or GF (*Tff3*) datasets. In addition, a number of key transcription factors involved in secretory cell and GC-specific fate determination (*Atoh1*, *Gfi1*, *Spdef*, *Foxa1*, *Foxa2*, *Foxa3*) were all positively correlated with postnatal age in ConvR but not GF mice, strongly indicating that microbiota colonization directs maturation of colonic GCs. Linked to this finding, functional analysis of age-correlated genes in all groups using the Reactome pathway database only identified significant pathway enrichments in the positively correlated ConvR dataset (**Fig. S1B**). This analysis highlighted microbiota-dependent induction of GC-enriched genes involved in COPI-mediated ER-Golgi anterograde protein transport (*Arf1, Arf2, Copa, Copg1, Copb2, Cope, Copz1*) and *O*-linked glycosylation genes involved in initialising (*Galnt4, Galnt6, Galnt10*), extending (*B3galt4, Gcnt3*), capping (*Fut2, Fut4, St6galnac2, St6galnac6*) or sulphating (*Chst4*) mucin *O*-glycan structures. Comparison of selected postnatal gene expression patterns indicated that a substantial degree of microbiota-dependent GC gene expression was associated with the neonatal-weaning transition, with several relevant genes (e.g. *Chst4, Gcnt3*) similarly regulated in both ConvR and GF tissues prior to weaning but diverging after weaning (**Fig. 3F**).

Lastly, we addressed the possibility that microbiota-dependent postnatal induction of GC-enriched gene expression was reflective of colonic GC numbers. It is frequently claimed that GF mice have fewer intestinal GCs than ConvR animals (*24*); however, the studies used to support this claim largely utilise non-specific histological staining of glycoprotein (e.g. lectins or Alcian Blue/Periodic Acid Schiff – AB/PAS) or do not specifically address the distal colon, which introduces a degree of uncertainty to this conclusion. To address this issue, we identified GCs using an intracellular marker by staining for the immature form of Muc2 (Apo-Muc2) that localises to the ER. Surprisingly, in adult tissues we identified a higher frequency of GCs in GF compared to ConvR colonic tissues (**Fig. 3G, H**). We went on to determine if microbiota-dependent divergence in GC numbers occurred in the postnatal period and found that in early postnatal tissues (P1-2) both GF and ConvR mice had an equally high GC frequency; however, at P3 and older the GC frequency in ConvR mice decreased towards the level observed in adult ConvR tissue (**Fig. 3I, S1C**). We confirmed decreased GC frequency at P3 by whole mount imaging of postnatal and adult tissues obtained from RedMUC2^98tr^ Muc2 reporter mice, which showed a clear epithelial transition between intercrypt regions with high GC frequency and epithelial cells emerging from developing crypts with lower GC frequency (**Fig. 3J**).

Together, these results demonstrated that microbiota colonization has a strong modulatory effect on postnatal colonic GC maturation. This modulation was characterized by reduction in total epithelial GC frequency in the early postnatal environment and subsequent induction of secretory transcription factors and gene programs associated with intracellular handling and modification (glycosylation) of secretory cargo proteins.

### Microbiota colonization drives postnatal maturation of colonic sentinel goblet cells

Having examined the role that the microbiota plays in guiding GC maturation, we next sought to determine if this maturation program included functional development of the sentinel goblet cell (senGC) GC subpopulation that acts as the secondary GC-intrinsic colonic defence mechanism. Colonic senGCs orchestrate inflammasome-dependent mucus secretion from upper crypt GCs in response to endocytosis of specific bacterial-derived molecules (microbe-associated molecular patterns; MAMPs) and we therefore first investigated dynamic *ex vivo* mucus growth rate changes in colonic tissues from GF mice and compared it to data from ConvR mice previously published by our group (*13*). We treated GF tissue with Carbachol (CCh, a cholinergic secretagogue) to quantify senGC-independent mucus secretion and with a range of bacterial MAMPs previously shown to drive variable senGC-dependent mucus secretory responses in ConvR tissue. GF tissue responded to both CCh and the bacterial MAMPs lipopolysaccharide (LPS), triacylated lipopeptide (P3CSK4) and Flagellin that drive senGC-dependent mucus secretion in ConvR tissues (**Fig. 4A**). Similarly to ConvR tissue, we did not observe induced mucus secretion when GF tissue treated with lipoteichoic acid (LTA), bacterial DNA, and the peptidoglycan subcomponents muramyl dipeptide (MDP) and g-D-glutamylmeso-diaminopimelic acid (iE-DAP) (**Fig. 4A**). Our data therefore indicated that MAMP-induced mucus secretion was conserved between GF and ConvR colonic tissues; however, when we compared the sensitivity of P3CSK4 induced mucus response in GF and ConvR tissues we found that GF tissues were more sensitive (∼36 fold) to P3CSK4 in comparison to ConvR tissues (**Fig. 4B**), indicating that GF and ConvR MAMP-induced secretory mechanisms may be divergent.

**Figure 4:**
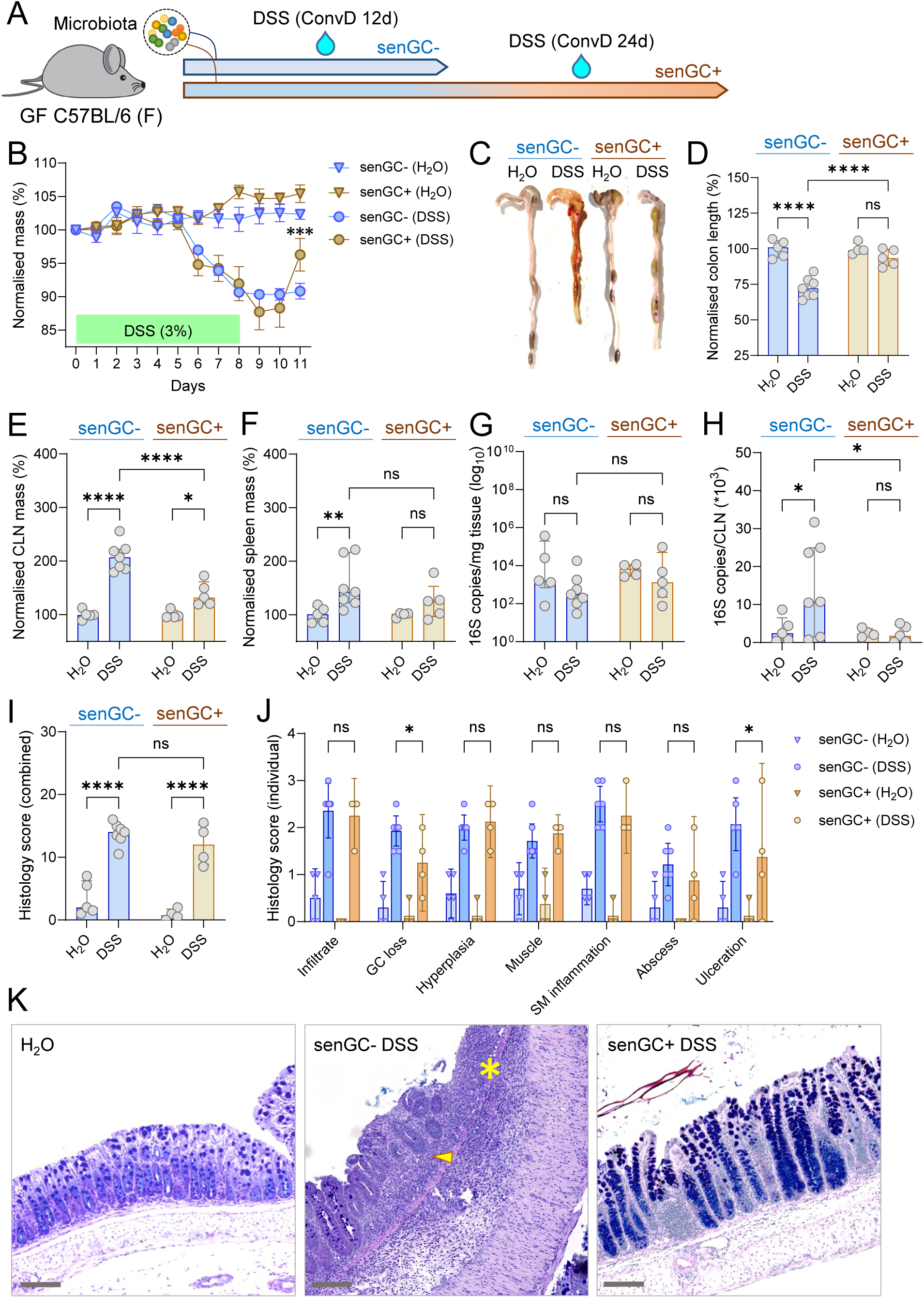
senGC maturation is microbiota-dependent. **A)** *Ex vivo* mucus growth in adult GF and ConvR mouse colon after stimulation with bacterial MAMPs. **B)** *Ex vivo* mucus growth dose-response to P3CSK4 in adult GF and ConvR mouse colon. **C)** *Ex vivo* mucus growth in adult GF and ConvR mouse colon stimulated with P3CSK4 in the presence or absence of senGC activation inhibitors. **D)** AB/PAS stained tissue sections from *ex vivo* experiments illustrated in (A). Emptied upper crypt GCs (red arrows) and lower crypt cavitation (yellow arrows) indicated. **E)** Whole mount confocal imaging of adult GF mouse colon treated with fluorescent Dextran tracer. Images show x/y-axis (upper panel) and x/z-axis (lower panel) cross-sections illustrating Dextran uptake by an upper crypt GC (purple arrow). **F)** *Ex vivo* mucus growth in neonatal (P3, 5, 15) and post weaning (P33) rat colon stimulated with P3CSK4 in the presence or absence of Dynasore inhibitor. **G)** Standardized expression of genes (columns) encoding known and predicted secreted proteins upregulated in mucus from P9-adult compared to P1-P7 rats (see Fig. 1H) in GC subpopulations (rows) identified by scRNA-seq. “Secretion” row indicates evidence of secretion determined by prior annotation or *in silico* prediction of classical or non-classical secretion by SecretomeP. **H)** Quantification of the frequency of Tgm3 expressing GCs as a proportion of the total GC population in neonatal (P3, P9, P14, P19) and post-weaning (P24) colonic tissue sections from ConvR mice. **I)** Confocal micrographs of representative tissue sections from P3 and P14 ConvR mice stained for Tgm3 (green) or the GC-binding lectins WGA (grey) and UEA1 (red). An individual GC from each image is indicated (yellow dashed line). Data represents n=3-5 (A-F, H-I)) animals per group, as indicated. All data is pooled from at least 2 independent litters or experiments. All histograms show median and interquartile range. Statistical comparisons between groups by 2-way ANOVA and Fishers LSD (A, C, F) or Kruskall Wallis and uncorrected Dunn’s test (H); p<0.05 (*), <0.01 (**) <0.001 (***), <0.0001 (****). Image scale bars are 50µm (D, I) or 20µm (E). # note: ConvR data displayed in A and B is reproduced from our previous publication (*13*) and is shown for illustrative purposes only.

To test whether GF MAMP-induced mucus secretion occurred *via* the senGC-dependent secretory pathway, we treated GF and ConvR tissues with P3CSK4 in the presence of inhibitors that block different elements of the senGC activation pathway. Inhibition of endocytosis (Dynasore), inflammasome activation (Caspase 1/11 inhibitory peptide) and MyD88 signalling (MyD88 inhibitory peptide) revealed that unlike ConvR tissues, P3CSK4 induced secretion in GF tissues was not diminished by endocytosis and inflammasome inhibitors, but remained dependent on TLR-MyD88 signalling (**Fig. 4C**). To visually assess the secretory response in GF tissues, we compared AB/PAS-stained sections of GF and ConvR tissues treated with P3CSK4 and CCh (**Fig. 4D**). In ConvR animals, we observed that P3CSK4 treated tissue had emptied GCs in the upper crypt region suggesting normal activation of senGCs, whereas CCh treatment resulted primarily in emptying of lower crypt GCs and crypt cavitation indicative of fluid secretion. Conversely, both P3CSK4 and CCh treatment of GF tissues induced GC emptying and cavitation in the lower crypt region, while upper crypt GCs were unaffected. Finally, we investigated if the lack of senGC-dependent secretion in GF tissues was the result of the absence of endocytotic GCs. We determined this by treating adult GF tissue with a fluorescent dextran tracer and imaging fixed whole mounts by confocal microscopy. Our results clearly demonstrate that GF upper crypts GCs successfully endocytosed fluorescent dextran (**Fig. 4E**). Together, these results suggest that GF colonic tissues mount a comparable MAMP-induced mucus secretion response to ConvR tissue; however, secretion in GF tissue does not follow a senGC-dependent secretory mechanism despite the presence of actively endocytosing upper crypt GCs.

Having determined that the senGC-dependent secretory response was microbiota regulated in adult mice, we next assessed its postnatal development in ConvR neonatal rat tissues in order to allow measurement of *ex vivo* senGC responses in neonatal samples. We examined *ex vivo* mucus growth in neonatal (P3, 5, 15) and post weaned (P33) rat colon stimulated with P3CSK4 in the presence or absence of Dynasore (**Fig. 4F**). Interestingly, the secretory response in P3 and P5 rats followed the senGC-independent secretory mode observed in adult GF mice, whereas this completely shifted to the senGC-dependent secretory response in P15 and P33 tissues, thus indicating an age dependent priming of senGCs in pre-weaned ConvR neonates.

Given that postnatal development of the senGC-dependent secretory response occurred over the same postnatal time frame where we had observed significant alterations in the colonic mucus proteome (**Fig. 1G, H**), we next sought to determine if there was any mucus proteomic evidence supporting senGC development. Prior single cell RNA sequencing (scRNA-seq) of colonic GCs has identified both a canonical and non-canonical GC differentiation trajectory, with expression of senGC activation pathway genes primarily localised to upper crypt non-canonical GCs (*23*) (**Fig. S2A**). We therefore screened expression of genes encoding known and *in silico* predicted secreted proteins upregulated in mucus in P9-adult compared to P1-P7 rats (**Fig. 1H**) in GC subpopulations identified by scRNA-seq. In this comparison, mucus protein enrichment over this postnatal period was significantly associated with higher standardized expression in upper crypt non-canonical GCs, notably Tgm3 and various proteases (Ctss, Ctsb, Ctsd, Mep1a) (**Fig. 4G, S2B**), thus identifying postnatal development of an upper-crypt non-canonical GC mucus proteomic signature. Since Tgm3 is a clear upper crypt non-canonical GC marker (**Fig. S2C**), we tracked the expression of this protein in colonic tissue sections from P3-P24 ConvR mice. We found that the frequency of Tgm3+ GCs increased from almost undetectable levels at P3 to a peak of ∼32% at P14, followed by a reduction to ∼22% post-weaning (**Fig. 4H, I**). Combined, this data demonstrated that microbiota-dependent postnatal maturation of the senGC-dependent secretory is associated with emergence of the upper crypt non-canonical GC lineage that has previously been linked to the senGC-dependent secretory response.

### senGC maturation requires complex microbiota colonization

In order to investigate the microbiota-dependent factors driving senGC maturation in an age-independent context, we attempted to induce senGC functional maturation in adult GF mice. Animals were conventionalized by faecal transfer from ConvR mice (ConvD) or monoassociated with the intestinal symbiont *Bacteroides fragilis*. Mice were sampled at 1-4 weeks post-colonization and assessed for the development of the senGC-dependent secretory response by *ex vivo* MAMP (P3CSK4) treatment (**Fig. 5A**). We detected development of the senGC-dependent response in 3w ConvD mice; however *B. fragilis* monoassociated mice failed to elaborate this response at any of the time-points tested. The lack of active senGC development in monoassociated mice could not be ascribed to lower colonization rates, as 16S qPCR analysis of stool DNA detected equally high bacterial load in monoassociated and ConvD samples (**Fig. 5B**). This indicated that a complex set of microbial signals, and not simply detection of common bacterial MAMPs by innate immune signalling mechanisms, are likely required to drive functional senGC maturation.

**Figure 5:**
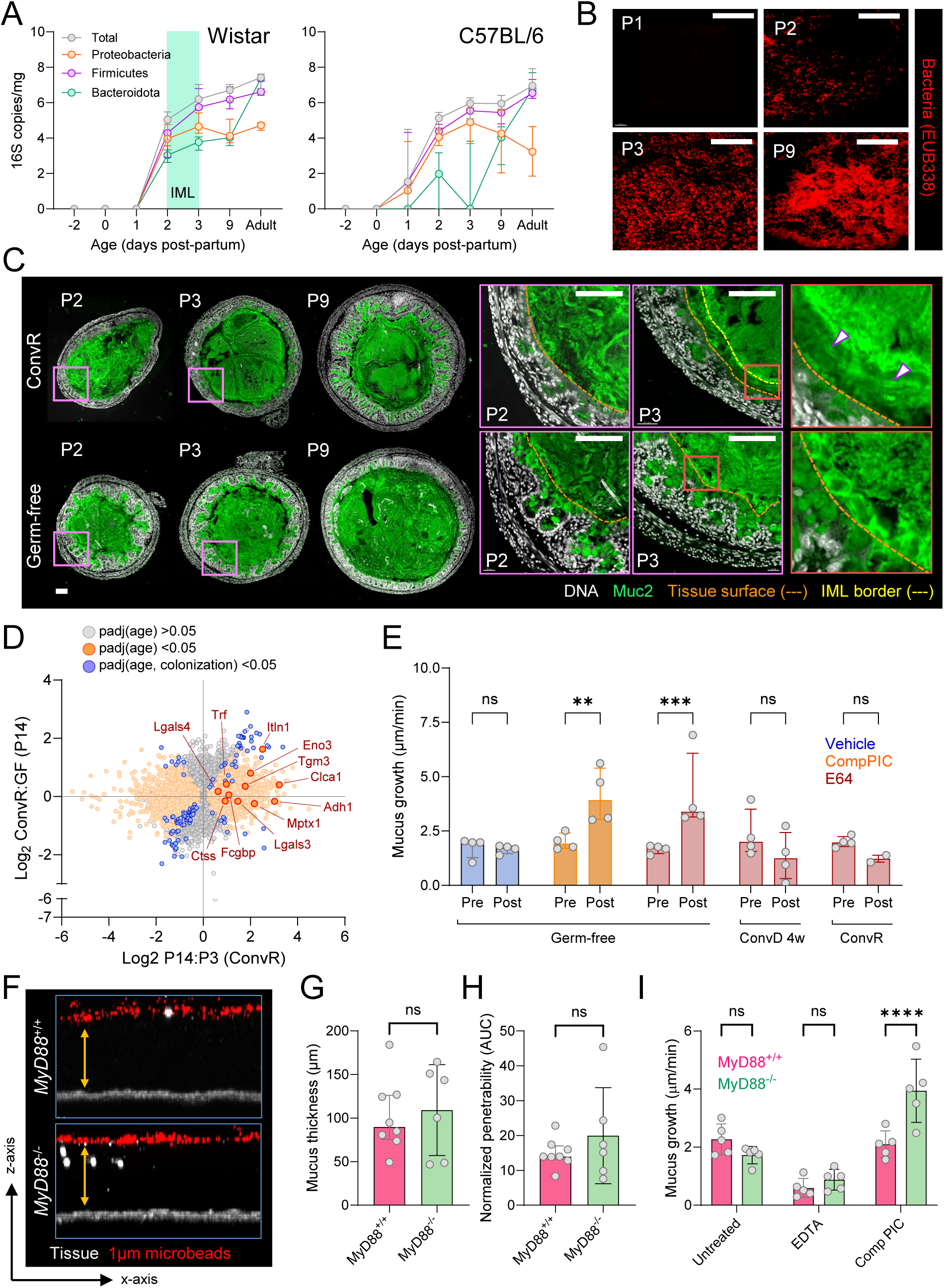
Microbiota-dependent induction of senGC function in adult GF mice. **A)** *Ex vivo* mucus growth in adult conventionalized (ConvD) and *B. fragilis* monoassociated mouse colon stimulated with P3CSK4 in the presence or absence of senGC activation inhibitors targeting endocytosis (Dynasore) or inflammasome activation (Casp IP). **B)** Total bacterial load in colon of conventionally raised (CR), ConvD and monoassociated mice quantified by 16S qPCR of stool DNA. Groups are colour-coded by the absence (senGC-; blue) or presence (senGC+; brown) of the senGC-dependent secretory response. **C)** *Ex vivo* mucus growth in adult *MyD88*^+/+^ and *MyD88*^-/-^ ConvR, GF and 4 week ConvD mouse colon stimulated with Flagellin in the presence or absence of a senGC activation inhibitor targeting inflammasome activation (Casp IP). **D)** *Ex vivo* mucus growth in adult *Nlrp6*^+/+^ and *Nlrp6*^-/-^ ConvR, GF and 4 week ConvD mouse colon stimulated with P3CKS4 in the presence or absence of a senGC activation inhibitor targeting inflammasome activation (Casp IP). **E)** Principle coordinate analysis of microbiota beta diversity (Bray-Curtis dissimilarity) based on metataxonomic 16S sequencing of DNA from ConvR and ConvD stool samples. Groups are colour-coded by the absence (senGC-; blue) or presence (senGC+; brown) of the senGC-dependent secretory response. **F)** Linear discriminant size effect analysis of bacterial taxa significantly enriched in stool from mice with the senGC-or senGC+ phenotype. **G)** Principle coordinate analysis of microbiota beta diversity (Bray-Curtis dissimilarity) based on metataxonomic 16S sequencing of DNA from ConvR WT, ConvD WT and ConvD *MyD88*^-/-^ and *Nlrp6*^-/-^ stool samples. Groups are colour-coded by the absence (senGC-; blue) or presence (senGC+; brown) of the senGC-dependent secretory response. **H)** Relative abundance of the genus *Mucispirillum* in ConvD *Nlrp6*^+/+^ and *Nlrp6*^-/-^ mice determined metataxonomic 16S sequencing of stool DNA. **I)** Standardized abundance (z-score) of bacterial taxa identified in (F) in 16S sequencing data from ConvD WT and ConvD *MyD88*^-/-^ and *Nlrp6*^-/-^ stool samples. Data represents n=4-7 animals per group, as indicated. All data is pooled from at least 2 independent experiments. All histograms show median and interquartile range. Statistical comparisons between groups by 2-way ANOVA and Fishers LSD (A, C, D), Kruskal Wallis and Dunn’s multiple comparison (B), PERMANOVA (E, G) or Mann-Whitney test (H); p<0.05 (*), <0.01 (**) <0.001 (***), <0.0001 (****).

In order to confirm that major innate immune signalling systems were not involved in senGC maturation, we conventionalised GF mice lacking either the TLR-MyD88 signalling axis (*MyD88*^-/-^) or the Nlrp6 inflammasome (*Nlrp6*^-/-^) and assessed the development of the senGC-dependent secretory response in 4w ConvD mice (**Fig. 5C, D**). Importantly, both MyD88 and Nlrp6 are involved in the senGC activation pathway, therefore requiring us to develop an approach to differentiate senGC-dependent and –independent mucus secretory responses in ConvD mice. Our prior data using MyD88 inhibition had established that TLR-MyD88 signalling was likely critical to both ConvR and GF mucus secretory responses (**Fig. 4C**). However, we have previously found that bacterial Flagellin can activate the senGC-dependent secretory pathway independently of MyD88 function (*13*). Accordingly, *ex vivo* mucus secretion was induced by Flagellin in ConvR *MyD88*^+/+^ and *MyD88*^-/-^ tissue in an inflammasome-dependent manner, whereas Flagellin drove inflammasome-independent secretion in GF *MyD88*^+/+^ tissue and had no effect on tissue from GF *MyD88*^-/-^ mice (**Fig. 5C**). Notably, Flagellin treatment of 4w ConvD tissue from both genotypes produced an identical response pattern to that observed in ConvR tissue (**Fig. 5C**), thus demonstrating that while MyD88 is a key regulator of both senGC-dependent ConvR mucus secretion, and senGC-independent GF mucus secretion, it does not appear to play a role in regulating microbiota driven development of functional senGCs. Conversely, application of a similar analytical approach to ConvR, GF and 4w ConvD *Nlrp6*^+/+^ and *Nlrp6*^-/-^ mice found that Nlrp6 function was only required for the senGC-dependent secretory response observed in ConvR mice; however, surprisingly, ConvD *Nlrp6*^-/-^ mice uniquely maintained the senGC-independent secretory response after colonization (**Fig. 5D**). While this result could be interpreted as suggesting that Nlrp6 is important for both the maturation and activation of senGCs, it should be noted that ConvR *Nlrp6*^-/-^ mice do not maintain senGC-independent secretory response. Consequently, this indicated that aspects of adult GF conventionalization that are divergent in *Nlrp6*^+/+^ and *Nlrp6*^-/-^ mice may play a role in the inability of ConvD *Nlrp6*^-/-^ animals to switch from the senGC-independent to the senGC-dependent MAMP-induced secretory response.

Previous studies have indicated that conventionalization of adult GF *Nlrp6*^-/-^ mice results in development of a microbiota configuration that is divergent from *Nlrp6*^+/+^ mice (*25*). Given that our data demonstrated that maturation of the senGC-dependent response was contingent on complex microbiota colonization, we hypothesised that variable senGC status in ConvD mice linked to differences in microbiota composition. To explore the composition of gut microbiota among various groups of ConvD mice, we employed 16S rRNA gene sequencing from extracted stool DNA. Beta diversity comparison using Bray-Curtis dissimilarity measures revealed significant differences in microbiota composition between groups lacking functional senGCs in early stages (1-2 weeks ConvD) and those with established senGCs function in later stages (3-4 weeks ConvD, ConvR donor mice; p=0.001) (**Fig. 5E**). We employed linear discriminant analysis effect size (LEfSe) analysis to identify specific bacterial taxonomic groups that associated with senGC status, which identified 6 taxa that were either positively (the genera *Muribaculaceae* and *Dubosiella* and family Prevotellaceae) or negatively (the genera *Bacteroides* and *Lachnospiraceae* and family Tanerellaceae) associated with functional senGCs (**Fig. 5F**). Subsequently, we determined the consistency of these observed differences by comparing the microbiota configurations between 4-week-old ConvD WT and MyD88^-/-^ mice (senGC positive) with those in 4-week-old ConvD Nlrp6^-/-^ mice (senGC negative). In this case, we did not detect any differences in overall microbiota composition between these groups (p>0.05) (**Fig. 5G**). Pairwise analysis of ConvD *Nlrp6*^+/+^ and *Nlpr6*^-/-^ mice did identify some differences in specific minor bacterial taxa, notably the genus *Mucispirillum* (**Fig. 5H**). However, none of the bacterial taxa that were correlated with senGC maturation in our previous analysis were altered between senGC positive or senGC negative mice under these conditions (**Fig. 5I**), thus indicating that such correlations were unlikely to be causal.

Combined, these results demonstrated that maturation of the senGC-dependent secretory response was inducible by full microbiota conventionalization of adult GF mice, and that this maturation could not be initiated by exposure to a single bacterium or by detection of bacterial MAMPs *via* MyD88 signalling.

### Microbiota-induced senGC maturation coincides with increased protection from colitogenic challenge

The senGC subpopulation is thought to play a role in preventing inflammation by protecting the colonic crypts from bacteria that breach the IML barrier. Postnatal, microbiota-dependent senGC maturation therefore presents an opportunity to study the impact of senGC proficiency or deficiency without the need for genetic modifications that risk off target effects. Furthermore, our ability to induce microbiota-dependent senGC maturation in adult GF mice in a colonization time-dependent fashion allows us to study this phenomenon independently of other factors involved in postnatal development, and to apply intestinal challenge models that cannot be applied to neonates.

Accordingly, we conventionalized adult GF mice for either 12 days (senGC-negative) or 24 days (senGC-positive) before exposing them to the colitogenic compound dextran sodium sulphate (DSS) in their drinking water (**Fig. 6A**). Mice were 3% DSS treated for 8 days, then switched back to normal water for 3 days recovery before sacrifice and tissue sampling. Daily monitoring of relative mouse mass found that both senGC-positive and -negative ConvD mice lost mass to a similar degree over the course of DSS treatment, with a significant difference only detected on the day of experimental termination that showed a more rapid recovery from colitis in senGC-positive mice (**Fig. 6B**). In addition, gross analysis of sampled colonic and lymphatic tissues found significant reductions in relative colon length (**Fig. 6C, D**) and increased relative mass of colon draining caudal lymph nodes (CLNs) and spleen tissue (**Fig. 6E, F**) in senGC-negative compared to the senGC-positive DSS-treated group. Quantification of tissue associated bacteria by 16S qPCR found a similar bacterial burden in the colonic mucosa of both senGC-negative and -positive groups (**Fig. 6G**); however, we detected higher bacterial load in CLNs isolated from the senGC-negative DSS-treated group (**Fig. 6H**). Lastly, we assessed colon tissue histopathology by scoring colonic swiss roll tissue sections for inflammatory infiltrate, GC loss, epithelial hyperplasia, muscle thickening, submucosal inflammation and the development of mucosal abscesses and ulceration (**Fig. 6I-K**). Overall, we did not observe any significant differences between the combined histology scores of the senGC-negative and senGC-positive DSS-treated groups (**Fig. 6I**); however, when analysing individual elements of the pathology index we noted significantly lower scores related to GC loss and ulceration in the senGC-positive DSS-treated mice compared to their senGC-negative equivalents (**Fig. 6J, K**).

**Figure 6:**
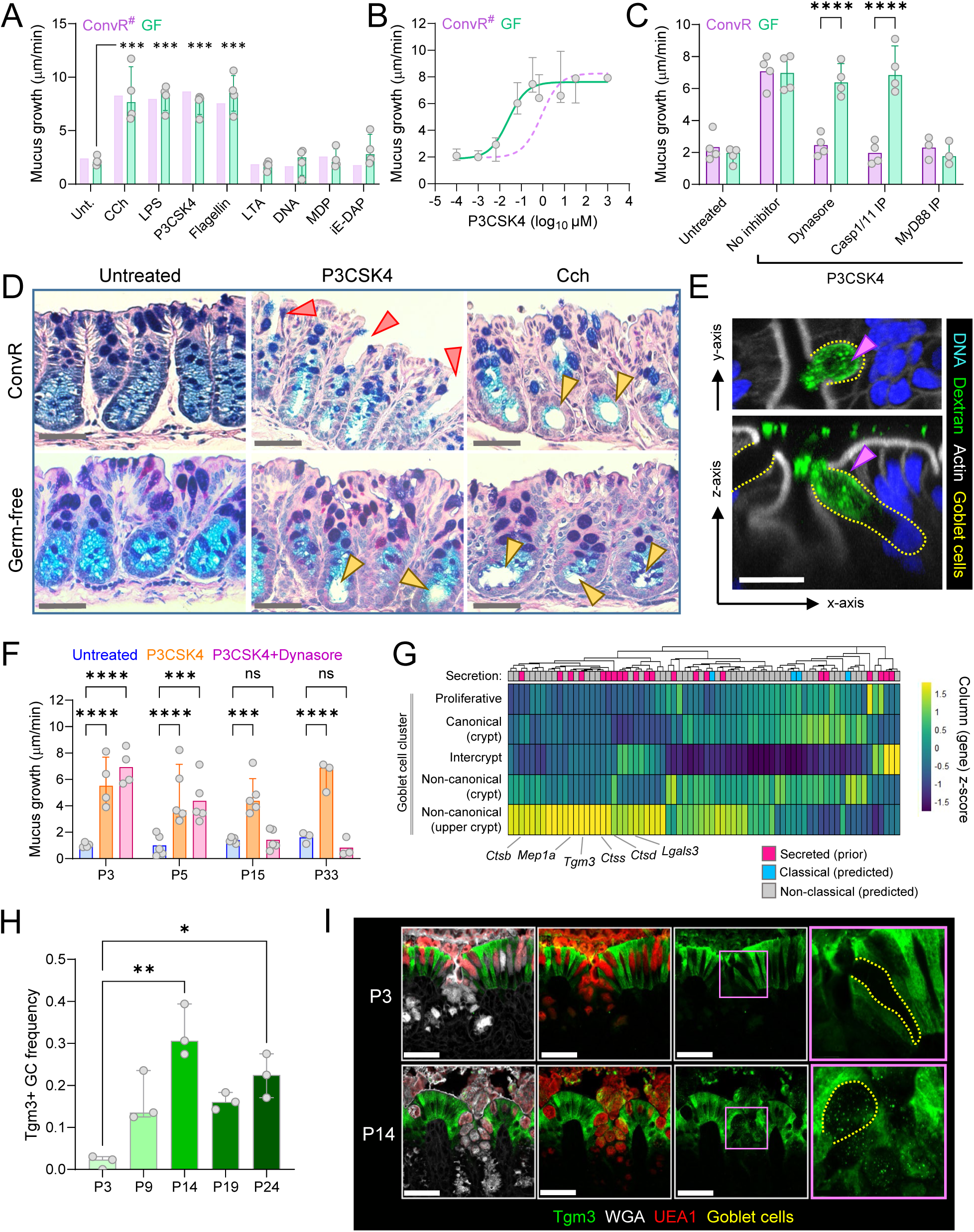
Induction of senGC maturation coincides with colitis protection. **A)** Timing of colitogenic DSS challenge in relation to microbiota-dependent induction of senGC maturation in adult GF mice. **B)** Tracking of day zero normalized mass changes in H_2_O and DSS treated ConvD mice from senGC- and senGC+ groups. **C)** Photographs of colonic tissue dissected from different treatment groups at termination of DSS challenge experiment. **D)** Quantification of total colon length in different treatment groups based on images shown in (C). All data normalized to average colon length of the H_2_O control animals for each group. **E)** Quantification of caudal lymph node (CLN) mass in different treatment groups. All data normalized to average CLN mass of the H_2_O control animals for each group. **F)** Quantification of spleen mass in different treatment groups. All data normalized to average spleen mass of the H_2_O control animals for each group. **G)** Determination of mucosal bacterial load in different treatment groups by 16S qPCR of colonic tissue DNA. **H)** Determination of bacterial translocation from the colon in different treatment groups by 16S qPCR of CLN DNA. **I)** Combined histology scores from different treatment groups determined by histopathological scoring of fixed colonic swiss roll tissue sections. **J)** Individual scores for different histopathology metrics from different treatment groups. **K)** Micrographs of AB-PAS stained fixed colonic swiss roll tissue sections used for histology scoring. Areas of ulceration (asterisk) and GC depletion (yellow arrow) in senGC-DSS treated tissue indicated. Data represents n=4-7 animals per group, as indicated. All data is pooled from 2 independent experiments. All histograms show median and interquartile range. Statistical comparisons between groups by 2-way ANOVA and Fishers LSD; p<0.05 (*), <0.01 (**) <0.001 (***), <0.0001 (****).

Consequently, these findings suggest that microbiota-dependent senGC induction coincides with increased protection from GC depletion and microbiota breach of the mucosal barrier during DSS-mediated colitis. Our experiments do not rule out that other factors may contribute to these phenotypes, thus it is not possible to definitively link a causal relationship to senGC maturation. However, given the proposed *in vivo* function of senGCs these findings may provide further support to this protective mechanism.

### Microbiota colonization matures senGC responses via regulation of Duox2

Previous studies of senGC activation in response to MAMPs indicate that a GC-intrinsic sequence of endocytosis, TLR-MyD88 signalling, ROS synthesis and Nlrp6 inflammasome assembly occur upstream of mucus secretion (**Fig. 7A**). Notably, analysis of RNA sequencing data from sorted GCs and colonocytes demonstrates that none of the confirmed or putative activation mediators are specifically expressed in GCs, on the contrary most show significantly higher expression levels in the colonocyte population (**Fig. 7B**). We therefore examined total bulk mRNA sequencing data from both 3w ConvD adult mice and P22 postnatal ConvR mice and compared them to their age-matched GF controls in order to establish if microbiota-induced expression of senGC pathway genes might indicate a common mechanism of senGC maturation (**Fig. 7C**). While microbiota-induced gene expression in both adult and postnatal mice did show a degree of overlap (notably induction of immune function-related genes), only 7.8% of induced genes and 6.4% of suppressed genes were significantly regulated in both groups (**Fig. 7D**). Indeed, most senGC pathway genes were either not significantly regulated by microbiota exposure, or were only induced in ConvD adult mice (*Nox1* and related accessory genes) or postnatal ConvR mice (*Nlrp6*) (**Fig. 7E**). The exception to this pattern was *Duox2* and its accessory *Duoxa2*, which were both strongly regulated by microbiota colonization in ConvD adult and ConvR neonate groups, thus indicating that expression of these genes might be linked to maturation of the senGC-dependent secretory response.

**Figure 7:**
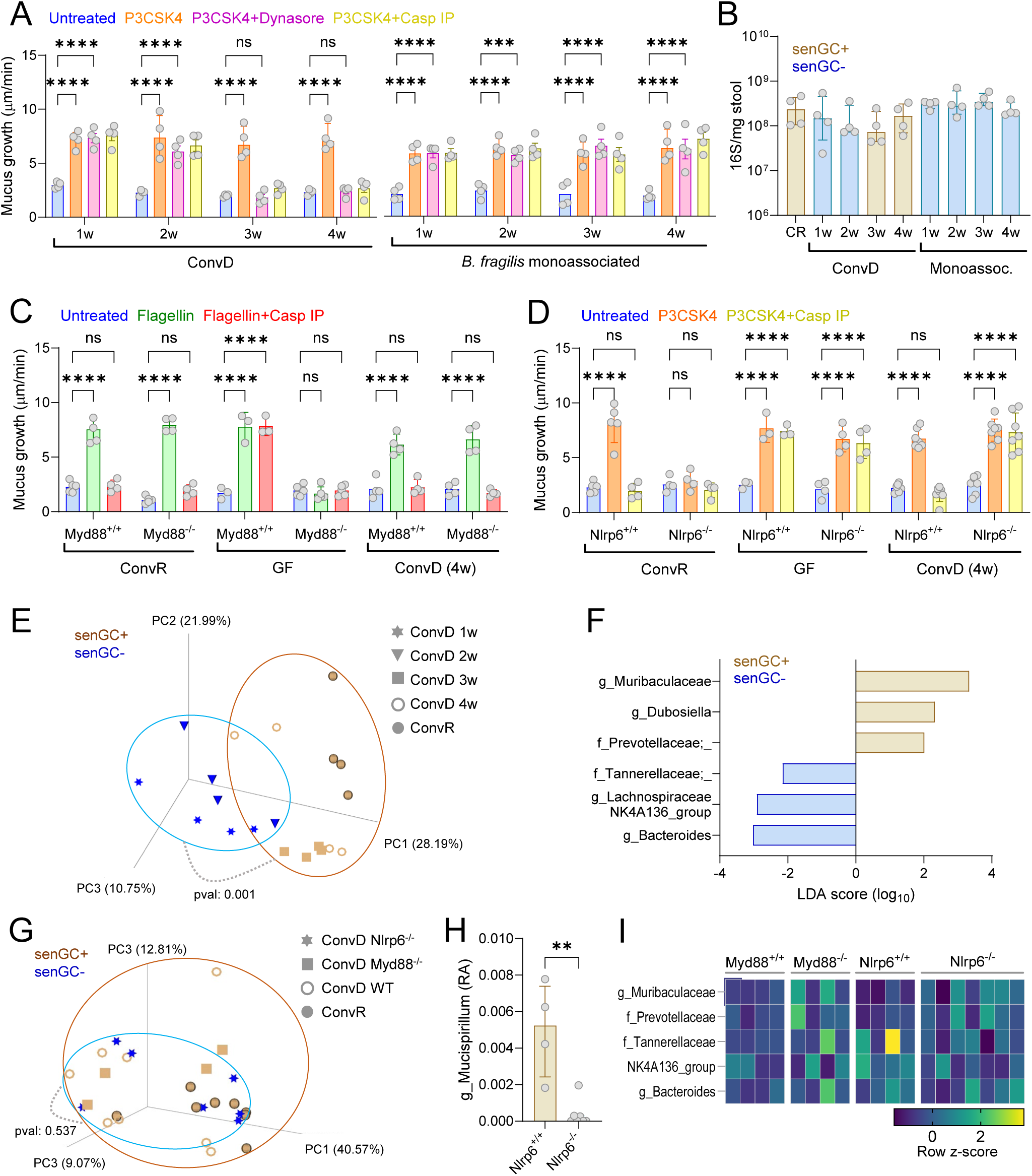
Microbiota induction of senGC maturation via regulation of Duox2. **A)** Schematic of the senGC activation pathway highlighting known (black) and putative (red) pathway genes. **B)** Expression of known and putative senGC genes in FACS isolated colonic GCs and colonocytes determined by DESeq2 analysis of bulk RNA sequencing data. **C)** Comparison of gene expression ratios between P22 ConvR:GF mice and adult 3-week ConvD:GF mice quantified by DESeq2 analysis of bulk colonic RNA sequencing data. Genes significantly up (red) or down (blue) regulated by microbiota exposure in both P22 and ConvD mice are indicated. **D)** Proportion of unique and shared genes significantly regulated by microbiota exposure in P22 ConvR and adult 3-week ConvD mice, based on data shown in (C). **E)** Comparison of microbiota-dependent expression of known and putative senGC activation pathway genes (A-B) in P22 ConvR and adult 3-week ConvD mice. Subset of data shown in (C). Genes not significantly regulated by microbiota in either group (grey), or genes regulated in either P22 ConvR (purple), adult ConvD (yellow) or both groups (teal) are indicated. **F)** Relative expression (compared to GF) of *Duox2* (left) and *Nox1* (right) genes in ConvD (brown) and *B. fragilis* monoassociated (blue) mice from 1-4 weeks colonization. Expression determined by qRT-PCR of colonic RNA, normalized to *Gapdh* and *Rplp0* expression. **G)** Expression of *Duox2* (left) and *Nox1* (right) genes in postnatal ConvR (purple; P3-33) and GF (teal; P9-P33) determined by DESeq2 analysis of bulk colonic RNA sequencing data. **H)** Confocal micrographs of fixed colonic tissue sections from ConvR WT mice stained for *Duox2* (left) and *Nox1* (right) mRNA by *in situ* RNA hybridization and counterstained by Epcam (grey). Duox2 or Nox1 expressing crypt regions indicated (yellow arrows). **I)** Confocal micrographs showing upper crypt GCs in fixed colonic tissue sections from ConvR, GF and ConvD mice stained for Duox2 (red), mucus (UEA1; green), Actin (grey) and DNA (blue). Duox2 or Nox1 expressing crypt regions indicated (yellow arrows). **J)** *Ex vivo* mucus growth in *Duox2*^fl/fl^ and *Duox2*^ΔIEC^ colon tissue treated with carbachol (CCh), LPS or P3CSK4. **K)** *Ex vivo* mucus growth in WT colon tissue treated with P3CSK4 in the presence or absence of the Nox1 inhibitor ML171. Data represents n=2-5 animals per group, as indicated. All data is pooled from at least 2 independent experiments or litters. All histograms show median and interquartile range. Statistical comparisons between groups by DESeq2 (B, C, E), Kruskal Wallis and Dunn’s multiple comparison (F) or 2-way ANOVA and Fishers LSD (J, K); p<0.05 (*), <0.01 (**) <0.001 (***), <0.0001 (****). Scale bars are 50µm (H) or 10µm (I).

We next sought to generate supportive evidence that regulation of Duox2 might provide a mechanistic basis for senGC maturation. Both Duox2 and its relative Nox1 are ROS generating NADPH oxidases that are expressed in the intestinal epithelium and have been previously linked to induction by the microbiota (*26–28*). Given that we had also observed *Nox1* induction in ConvD adult mice, and that both enzymes could theoretically be involved in the senGC activation pathway, we therefore characterised induction of both Duox2 and Nox1 over the course of our different colonization models. RNA expression of *Duox2* and *Nox1* by qPCR analysis of both ConvD and *B. fragilis* monoassociated tissue demonstrated that both genes were significantly induced in the ConvD, senGC positive colonization experiment, but were only weakly or not induced in the monoassociated mice that failed to develop an active senGC response (**Fig. 7F**). Conversely, analysis of ConvR (P3-P33) and GF (P9-P33) mRNA sequencing data demonstrated microbiota-dependent induction of *Duox2* but not *Nox1* in postnatal tissues (**Fig. 7G**).

As functional senGCs have only been observed in the upper crypt epithelium, we next spatially characterized expression of both genes by RNA *in situ* hybridisation in mouse colonic tissue sections (**Fig. 7H**). In line with previous observations (*27*), *Nox1* expression was restricted to the crypt base; however, we observed *Duox2* expression at both the crypt base and upper crypt regions. Furthermore, we further validated Duox2 expression in upper crypt GCs by immunostaining and observed an expression pattern indicating that in ConvR mice Duox2 was localised to a continuous subcellular compartment that likely represents the endoplasmic reticulum as it did not co-localise with glycosylated Muc2 stained by UEA1 (**Fig. 7I**). Importantly, this Duox2 staining pattern was present, but notably more fragmented and weaker in tissue from GF mice but similar to ConvR mice in tissue from ConvD adult mice. Consequently, our expression and localisation data supported a potential role for Duox2, but not Nox1, in regulating senGC activation.

Accordingly, we investigated the mechanistic requirement for Duox2 in senGC activation by *ex vivo* analysis of the senGC-dependent secretory response in mice deficient for intestinal epithelial Duox2 expression. We generated constitutive conditional knockout mice by crossing *Duox2*^fl/fl^ with Vil1-Cre mice to generate *Duox2*^ΔIEC^ offspring and quantified inducible mucus secretory responses. Both *Duox2*^fl/fl^ and *Duox2*^ΔIEC^ littermates had similar baseline mucus growth rates and secretion in response to the cholinergic agonist carbachol; however, *Duox2*^ΔIEC^ lacked the MAMP (LPS, P3CSK4) induced senGC-dependent response (**Fig. 7J**), demonstrating that Duox2 is functionally critical for senGC activation. Conversely treatment of WT tissue with the Nox1 specific inhibitor ML171 had no impact on MAMP-induced secretion, further indicating that Nox1 is unlikely to be directly involved in this pathway (**Fig. 7K**).

Consequently, our data shows that microbiota-dependent induction of Duox2 expression is a common feature of postnatal and adult mice that go on to develop mature senGCs. Given that our subsequent analysis of Duox2 function demonstrates that this enzyme is a key mediator of senGC activation, our data indicates that microbiota induction of Duox2 expression in GCs is an important element of postnatal senGC maturation.

## Discussion

In this investigation we have defined the postnatal maturation of the colonic IML and senGC protective mechanisms in mixed rodent model systems. We establish that these GC-dependent protective functions develop in sequence in the early pre-weaning environment, with IML barrier function emerging almost immediately *post-partum* and IML post-secretory processing dynamics and the senGC response developing over the subsequent postnatal period. We further establish that different aspects of this development and GC maturation are regulated either independently or dependently on microbial colonization, with mechanistic roles for MyD88 signalling in modulating post-natal IML proteolytic processing, and a causal link between microbiota induction of Duox2 expression and priming of the senGC-dependent secretory response.

As previously established in adult GF mice (*12, 18*), we have demonstrated that postnatal development of IML barrier function and faecal encapsulation of the microbiota by Muc2 occurred in a microbiota-dependent fashion that was independent of classical innate immune signalling involving MyD88. Along with previous findings that immune signalling *via* the inflammasome is not required for baseline IML formation (*12, 29*), these results further support the concept that microbiota-dependent maturation of this system occurs independently of innate immunity. While our current data does not suggest any candidate regulatory mechanisms, the rapidity of postnatal IML maturation over the first three days *post*-*partum* implies that this system develops in response to a well conserved and fast-acting signal(s), which likely rules out microbial metabolites (e.g. fermentation products) that would take time to build up in the intestinal environment. A possibility that we have not addressed in this investigation is that the colonization status of the mother, rather than incoming microbial colonizers themselves, regulates IML maturation. Prior analysis of small intestinal tissues from GF pups littered by gestationally colonized GF mothers indicates maternal regulation of mucus modulatory ion channel and mucus component genes including Clca1 and Zg16 (*30*). Examination of colonic IML development in a similar system is a promising future avenue of research.

The postnatal dynamics of IML modulatory factors illustrates a highly mixed landscape of microbiota-dependent and independent regulation. Critical shapers of Muc2 polymeric network integrity (Tgm3) (*7*) and metalloprotease-dependent mucus expansion (Clca1) (*11*), while strongly age regulated at both the mRNA and protein abundance level, were not regulated by colonization status in the pre-weaning environment. However, our data did point to the existence of cysteine protease-mediated suppression of mucus expansion in the immediate postnatal period that was regulated by microbiota colonization *via* MyD88 signalling. This indicates that the reduced mucus growth observed in this period is partly the result of active suppression by the colonic tissue, and it is tempting to speculate that this may reflect a mechanism to slow down mucus turnover during the period that the IML barrier is being established; however, this phenomenon requires further investigation. We furthermore identified microbiota-dependent modulation (both induction and repression) of multiple genes involved in the Muc2 *O*-glycosylation machinery and GC-fate targeted transcription factors, that were especially regulated in the immediate pre- and post-weaning time period. These findings point to a dynamic and coordinated process of postnatal GC and IML maturation that likely reflects the physiological needs of mobilising intestinal content during the initial switch from *in utero* to enteral nutrition, and subsequently adapting to the significantly increased microbial load and diversity associated with the weaning transition (*31*).

Development of the bacterial MAMP-induced senGC mucus secretory response was observed over the first 1-2 postnatal weeks. Intriguingly, both early neonatal and adult GF colonic tissue responded to MAMPs in a senGC-independent fashion that maintained functional reliance on intact MyD88 signalling but was independent of the active endocytosis and Nlrp6 inflammasome activation required for senGC activation. Histological analysis indicated that senGC-independent responses were more likely the result of fluid rather than mucus secretion, and we speculate that this may function as a rapid response mechanism to flush colonizing bacteria out of developing epithelial crypt structures in the early colon. Intriguingly, the timing of senGC functional maturation was coincidental with postnatal changes in IML properties and increased abundance of key mucus regulatory factors (e.g. Tgm3) that we had previously observed. Our prior work has linked senGCs to a subpopulation of colonic GCs referred to as non-canonical GCs, due to their expression of genes that are normally enriched in non-secretory epithelial cells. Examination of mucus proteins with increased abundance over the senGC development window identified a distinct non-canonical GC signature in the mucus proteome, including proteins with known (Tgm3, Lgals3) or putative (Mep1a, Ctsb, Ctsd, Ctss) roles in regulating mucus protective functions (*7, 10*). Consequently, our data indicates that senGCs and other non-canonical GCs develop in the preweaning environment and may have a direct impact on IML maturation *via* the production and secretion of mucus regulatory factors that are not expressed by canonical GCs.

Finally, our data demonstrated that postnatal development of functional senGC secretion was driven by microbiota colonization and was likely to be mechanistically driven by induction of Duox2 expression that primes the senGC activation pathway. Previous work has established microbiota regulation of Duox2 expression (*26*) and linked H_2_O_2_ production by Duox2 to activation of TLR signalling (*28*) and we now establish a functional role for Duox2 in the senGC activation pathway. Duox2 protein expression in upper crypt GCs was localised to an intracellular structure likely corresponding to the ER or Golgi compartment. Previous work investigating GC uptake of luminal material has found that endocytotic cargo can be trafficked to either of these organelles (*14*), and this Duox2 localisation aligns well with our previous work establishing that ROS synthesis functions downstream of endocytosis and TLR signalling in the senGC activation pathway (*13*).

The postnatal period represents a crucial window of time where the intestinal mucosa must quickly adapt to both tolerate and defend itself from colonizing microorganisms (*32*). Interruption of this process prevents the tolerogenic imprinting of the mucosal immune system that underlies intestinal homeostasis (*17, 33*), resulting in increased susceptibility to inflammatory disease in later life. GC-intrinsic protective functions are critical legislators of healthy intestinal microbiota-host interactions, and the developmental stages that we have described in this investigation are highly likely to be important aspects of neonatal, infant and adult health.

### Study Limitations

Our investigation has employed widely used rodent (both mouse and rat) experimental models to define developmental stages of GC-intrinsic protective maturation. However, it should be emphasised that the timing of human intestinal maturation is different to rodents, with significantly more development (e.g. colonic epithelial crypt formation) occurring during gestation in humans. Because of this divergence, it is possible that the microbiota-independent developmental dynamics we have observed in rodents (e.g. increased Tgm3 or Clca1 mucus abundance) may occur in the prenatal stage in humans. Furthermore, as discussed previously we have not determined whether microbiota-dependent developmental dynamics are driven directly by postnatal colonization of the neonate, or by exposure to maternally derived microbial products *in utero*. If such dynamics are related to maternal microbial status, it is possible that these too could occur at the prenatal stage. Lastly, while our work has revealed many dynamic mucus and GC phenomena and has established a range of causal mechanistic associations, many aspects remain undetermined. These include the identity of the cysteine protease that appears to suppress IML growth during the early neonatal period, the physiological relevance of this suppression or the precise microbial modulators of various microbiota-dependent phenomena (e.g. senGC activation). However, we believe that our findings provide a wide-ranging foundation for the further study of these phenomena and highlight the importance and experimental utility of studying the pre-weaning period for understanding fundamental mucosal biology.

## Acknowledgements

We gratefully acknowledge the support of staff at the Genomics and Bioinformatics, Experimental Biomedicine and Centre for Cellular Imaging Core Facility platforms at Sahlgrenska Academy, University of Gothenburg. MVS and GMHB are supported by funding from Knut & Alice Wallenberg Foundation *via* the Wallenberg Centre for Molecular & Translational Medicine at the University of Gothenburg. GMHB is further supported by grants from Vetenskåpsrådet (2018–02278, 2023–02283), Cancerfonden (210301-JIA, 211422-Pj), IngaBritt & Arne Lundbergs Foundation (LU2022-0069) and the National Institute of Allergy and Infectious Diseases (5U01AI095542-10). FS and PR are supported by the Deutsche Forschungsgemeinschaft (DFG) Research Unit 5042 “miTarget” (TP3 P.R. and TP5 F.S.), Collaborative Research Center (CRC) 1182 “Origin and Function of Metaorganisms” (subprojects C02, P.R.) and the Cluster of Excellence Precision Medicine in Chronic Inflammation. We would like to sincerely thank all past and present members of the Mucin Biology Groups research cluster for all discussions and suggestions relating to the development of the project.

## Author contributions

Conceptualisation: GMHB; Study design and methodology: AJ, MVS, BY, EN, EL, LA, FS, PR, LV, LMH, AW, TP, MEVJ, GMHB; Investigation & Analysis: AJ, MVS, BY, EN, EL, LA, GMHB; Writing – Original Draft: MVS, GMHB; Writing – Review & Editing: AJ, MVS, BY, EN, EL, LA, FS, PR, LV, LMH, AW, TP, MEVJ; Visualisation: GMHB; Supervision & Funding: MEVJ, GMHB.

## Methods

### Experimental animals

All animals used in this study were housed under specific pathogen free conditions with *ad libitum* access to food and water with a 12h light/dark cycle. All mice were on a C57BL/6N background and bred in-house. Both non-pregnant and timed-mated Wistar rats were purchased from Charles River. Germ-free (GF) mice were maintained in sterile flexible film isolators and monitored regularly by aerobic and anaerobic culturing as well as by PCR for bacterial 16S rRNA. The generation of RedMUC2^98tr^ Muc2 reporter mice, *MyD88^-/-^*, *Nlrp6^-/-^* and *Duox2* intestinal epithelium conditional knockout mice has been previously described (*13, 34–36*). Experimental groups consisted of neonatal, weaned and adult (12-20 week-old) mice and rats as indicated for each experiment. All animals examined were either female only, or a balanced mixture of males and females. All experimental groups compared in the study were fed identical rodent chow diets and all comparisons of wild-type and null mutant knockout mice used co-housed littermates generated by breeding heterozygous animals. For full conventionalisation of GF mice, colonic stool and caecal content was pooled from three conventionally raised (ConvR) donor mice and homogenized in PBS supplemented with 1% (w/v) cysteine and immediately gavaged (200 µl) into GF recipient mice. Conventionalised mice were subsequently housed under normal ConvR conditions. For monoassociation of GF mice with *Bacteroides fragilis*, strain NCTC 9343 was grown at 37°C under anaerobic conditions in BHI broth supplemented with 0.05% (w/v) cysteine and 100 nM Hemin, and 200 µl of overnight culture was gavaged into GF recipient mice. Monoassociated mice were subsequently housed in sterile isolator cages. All weaned and adult animals were anesthetized using isoflurane and killed by cervical dislocation before collection of samples. All neonatal animals were killed by decapitation. All experimental procedures involving animals were approved by the Swedish Laboratory Animal Ethical Committee in Gothenburg.

### Ex vivo analysis of colonic mucus properties

Colonic mucus thickness, growth rate and barrier function were determined by *ex vivo* analysis of live colonic tissue as previously described (*29, 37, 38*). Briefly, fresh tissues were collected by dissection and flushed using cold oxygenated Krebs buffer. Tissues were opened longitudinally, the muscle layer was removed by microdissection and then mounted mucosa-side up in a custom-made horizontal perfusion chamber with basolateral perfusion of oxygenated Krebs-glucose buffer (10 mM), static apical oxygenated Krebs-mannitol (10 mM) and heated to 37°C. For mucus thickness and growth rate measurements, the mucus surface was visualised using 10 µm black microbeads (Polybead) diluted 1:20 in Krebs-mannitol. Mucus and tissue were observed using a stereomicroscope and the mucus thickness measured using a 5 µm micropipette linked to a micrometer device (Mitotoyo). Mucus thickness was determined at t=0 min, and mucus growth rate was determined by measuring mucus thickness at t=30 min and t=60 min. In some experiments tissues were treated with either EDTA (5 mM; Merck), 1x complete EDTA-free protease inhibitor cocktail (Roche) or E64 (20 µM; Merck) inhibitors at t=30 min. For mucus barrier function measurements, tissues were mounted in confocal microscope adapted imaging chambers without perfusion or heating. Mucus was overlaid with a Krebs buffer solution containing Syto9 tissue dye (12.5 µM; ThermoFisher) and 1 µm crimson Fluospheres (1:20; ThermoFisher) for 5 min, after which excess dye and beads were removed by washing with Krebs-mannitol. Tissue and Fluospheres were imaged using an LSM700 laser scanning confocal microscope equipped with an x20 water-immersion objective, 488/639-nm lasers, and Zen acquisition software (Carl Zeiss). Fluorescent signals were mapped using Imaris software (Oxford Instruments) z-axis positions of tissue and Fluospheres was extracted. Mucus barrier function (normalised penetrability) was quantified by analysis of Fluosphere distribution within the mucus layer. Frequency distribution curves were generated for each z-stack using Prism 9 software (GraphPad) normalized to maximum frequency values and then normalized to the position of the mucus surface. Area under the curve (AUC) data expressed as normalized penetrability was used for quantitative comparison of Fluosphere penetration into the mucus layers of different samples.

### MAMP-induced mucus secretion

The capacity of colonic tissues to secrete mucus in response to bacterial MAMPs was performed using the same *ex vivo* experimental setup as described above for measuring mucus growth rates with a micropipette mounted micrometer. All bacterial MAMPs were purchased from Invivogen and were synthetic or Ultrapure grade. Mucus growth was measured at t= 0 and t= 30min to establish baseline growth rates. At t= 30min, tissues were treated apically with LPS from *E. coli* O111:B4 (200 µg/ml), LTA from *S. aureus* (200 µg/ml), P3CSK4 (50 µg/ml), DNA from *E. coli* (200 µg/ml), flagellin from *B. subtilis* (50 µg/ml), MDP (200 µg/ml), iEDAP (200 µg/ml) or basolaterally with Carbachol (Merck; 1 mM). Mucus growth was measured at t= 45min (15 min post treatment) to determine the impact of treatments on growth rate. All MAMPs and Carbachol were reconstituted in ddH_2_O, sonicated and stored at −20°C until use. For experiments using inhibitors to block MAMP-induced mucus secretion, inhibitory compounds were added to apical buffer solutions at t= 0 and maintained throughout the experiment. Inhibitors used were Dynasore (Merck; 100 µM), Ac-YVAD-cmk Caspase 1/11 inhibitory peptide (Merck; 100 µM) and Pepinh-MYD MyD88 inhibitory peptide (Invivogen; 100 µM).

### Mass spectrometry-based profiling of the mucus proteome

Samples were collected *ex vivo* from distal colonic tissues mounted in horizontal perfusion chambers as described above. Mucus was aspirated form the mucosal surface using Maximum Recovery pipette tips (Axygen), mixed with 2x cOmplete protease inhibitor cocktail (Merck) and stored at −80°C until analysis. Sample processing was performed as previously described (*38*). Briefly, mucus was reduced overnight in 6 M guanidinum hydrochloride, 0.1 M Tris/HCl (pH 8.5), 5 mM EDTA, 0.1 M DTT (Merck) followed by filter aided sample preparation adapted from a previously developed protocol (*39*) using 10 kDa cut-off filters (Pall Life Sciences). Proteins were alkylated with iodoacetamide (Merck) and sequentially digested on the filter with LysC (Wako) and trypsin (Promega). Peptides were cleaned with StageTip C18 columns prior to MS analysis.53 NanoLC–MS/MS was performed on an EASY-nLC 1000 system (ThermoFisher), connected to a QExactive Hybrid Quadrupole-Orbitrap Mass Spectrometer (ThermoFisher) via a nanoelectrospray ion source. Peptides were separated using an in-house packed reverse-phase C18 column with a 60-min 4–32% acetonitrile gradient. Mass spectra were acquired from 320–1,600 m/z at resolution 70,000, and the 12 peaks with highest intensity were fragmented to acquire the tandem mass spectrum with a resolution of 35,000 and using automatic dynamic exclusion.

Proteins were identified using MaxQuant (v1.5.7.4) (*40*) searching the mouse UniProt protein database supplemented with mouse mucin sequences (http://www.medkem.gu.se/mucinbiology/databases/). Searches used full tryptic specificity, maximum 2 missed cleavages, 20 ppm precursor tolerance for recalibration search followed by 7 ppm for the final search, and 0.5 Da for fragment ions. Modifications were set as carbamidomethylation of cysteine (fixed), methionine oxidation (variable) and protein N-terminal (variable). The FDR was set to 1% both for peptide and protein levels and minimum peptide length was set to 6 amino acids. Proteins were quantified using label-free quantification (LFQ) using at least two peptides for quantification. LFQ data was analyzed using Perseus (v1.6.2.2) (*41*). Proteins were filtered for potential contaminants and detection in at least 50% of samples from one experimental age group. Data was log10 transformed and missing values were imputed from a normal distribution using default settings. Two-sample Welch’s t test with Benjamini-Hochberg FDR were used to identify specific protein abundance differences between experimental groups. Principal component analysis (PCA) was used to visualise clustering of different sample groups.

For certain analyses, proteins identified by mass spectrometry were defined as “secreted” based on prior knowledge, database annotations or *in silico* prediction. Individual protein localisation annotations and amino acid sequences were retrieved from UniProt. For *in silico* predication of secretion, protein amino acid sequences were analysed using SecretomeP (v 2.0) (*42*). Predicted classical secretion was defined as the presence of a signal peptide sequence, predicted non-classical secretion was defined as the absence of a signal peptide sequence and an NN-score >0.6.

### Histology

Colonic tissues were fixed in either methanol-Carnoy (Figures 1E, 2C, 3G) or formalin (Figures 4I, 6K, 7I) solutions for at least 24 h, and tissue was subsequently paraffin embedded and cut into 5 μm thick transverse sections. Tissue sections were deparaffinised by sequential washing in xylene substitute (20 min at 60°C; Merck) and 100% (5 min), 95% (5 min), 70% (5 min), and 30% (5 min) ethanol. For histochemical staining, tissue sections were stained with Alcian blue and Periodic acid-Schiff (AB/PAS) stains as previously described (*5*). For fluorescent staining, antigen retrieval was performed by immersion of sections in 10 mM citrate buffer (95°C, 30 min). Sections were washed in PBS, permeabilized for 5 min using 0.1% vol/vol Triton X-100 (Merck), and blocked using 5% vol/vol FCS. Mature Muc2 and Apo-Muc2 were detected using in-house Muc2C3 (*5*) and PH497 (*43*) rabbit polyclonal primary antibodies respectively. Tgm3 was detected using NBP1-57678 (Novus Biologicals) rabbit polyclonal primary antibody. Duox2 was detected using anti-Duox1/2 I2 (*44*) rabbit polyclonal primary antibody kindly provided by Professor Francoise Miot (Université libre de Bruxelles). Actin was detected using MAB1501 (Merck) mouse monoclonal primary antibody. Sections were incubated with primary antibodies overnight at 4°C and subsequently washed in PBS and stained with goat anti-rabbit Alexa 488 or Alexa 555-conjugated secondary antibodies, or goat anti-mouse Alexa 647-conjugated secondary antibody (ThermoFisher) for 2 h at room temperature. Lastly, slides were washed with PBS and counterstained with a Hoechst-34580 DNA dye (5 μg/mL; Merck) in some cases supplemented with combinations of UEA1 Atto 488-conjugated lectin (10 µg/mL; Merck), UEA1 DyLight647-conjugated lectin (10 µg/mL; Vectorlabs) or WGA Alexa 555-conjugated lectin (10 µg/mL; ThermoFisher) for 15 min. Slides were rinsed in dH2O, coverslipped using ProLong Gold Antifade mountant (ThermoFisher) and imaged using an LSM700 confocal microscope (Zeiss) with a x20 air objective.

### In situ hybridisation

Fluorescence *in situ* hybridisation (FISH) was used to detect bacterial cells and specific mRNA transcripts in histological tissue sections. FISH staining for bacterial 16S rRNA was performed using methanol-Carnoy fixed tissue sections described above. Sections were deparaffinized by sequential washing in Xylene substitute (20 min at 60°C; Merck), 100% ethanol (5 min), and 95% ethanol (5 min). Slides were air dried and flooded with hybridization buffer (40% v/v formamide, 0.1% w/v SDS, 0.9 M NaCl, and 20 mM Tris, pH 7.4) supplemented with Alexa 555–labelled universal bacterial FISH probe EUB33840 (1 mM). Slides were incubated at 37°C overnight in a RapidFISH Slide Hybridization Oven (Boekel Scientific), subsequently submerged in wash buffer (0.9 M NaCl and 25 mM Tris, pH 7.4), and incubated for 20 min at 50°C. Lastly, slides were rinsed in double-distilled water and counterstained with Hoechst dye (5 μg/mL; Merck). Stained slides were subsequently imaged using an LSM700 confocal microscope (Zeiss).

FISH staining for mRNA transcripts was performed using the RNAscope technique in combination with formalin fixed tissue sections. Tissue sections were processed for FISH staining using the RNAscope Multiplex Fluorescent Reagent Kit v2 (Advanced Cell Diagnostics) in combination with probes for either *Duox2* (Mm-Duox2) or *Nox1* (Mm-Nox1) transcripts as per manufacturers instructions. After hybridisation, sections were washed 3 times with 10% v/v Tween20-TBS and blocked with 1% w/v BSA in Tween20-TBS for 20 min.

Sections were counterstained for Epcam by overnight incubation at 4°C with HPA026761 (Merck) rabbit primary antibody. Sections were washed in Tween20-TBS, and primary antibody was detected by incubation with goat anti-rabbit Alexa Fluor 647 secondary antibodies (ThermoFisher) for 2 h at room temperature. Stained slides were subsequently imaged using an LSM700 confocal microscope (Zeiss).

### Endocytosis and tissue whole mount imaging

Colonic tissue was mounted in horizontal perfusion chambers as described above and treated apically with Dextran Alexa 488-conjugated tracer (ThermoFisher) for 15 min. Excess tracer was washed away and the tissue fixed in formalin for 1 h and then permeabilized with 0.5% (v/v) Triton X-100 for 15 min and washed with PBS. Tissue was stained for 1 h with a mixture of Hoechst-34580 DNA dye (10 µg/mL; Merck) and Phalloidin Alexa 647 conjugate (1:400; ThermoFisher) to visualize actin. Stained tissue was washed with PBS, transferred to a microscope slide and coverslipped with Prolong-Gold Antifade mounting medium (ThermoFisher). Whole mounts were visualized by generating confocal z-stacks using an LSM 700 microscope (Zeiss) with a x40 oil immersion objective.

### RNA extraction, qRT-PCR and sequencing

Distal colonic tissue was fixed for RNA preservation using RNAlater solution (Qiagen). Tissues were lysed in RLT buffer using an Ultra-Turrax rotor-stator homogenizer (IKA Werke) and RNA extracted using RNeasy Mini columns (Qiagen) and eluted into RNase-free H_2_O according to the manufacturers instructions. The quality of isolated RNA was determined using an Experion Automated Electrophoresis platform (Bio-Rad), and samples were stored at −80°C until further analysis.

Expression of *Nox1* and *Duox2* genes were analysed by qRT-PCR of cDNA prepared from 600 ng RNA extractions using the High-Capacity cDNA Reverse Transcription Kit (ThermoFisher). PCRs (20 µl) were prepared using SsoFast EvaGreen Supermix (Bio-Rad), 450 nM forward and reverse primers, and 10 ng cDNA. PCR cycling conditions were 95°C for 2 min and 40× cycles of 95°C for 5 s and 58°C for 30 s. Reactions were monitored using a CFX96 platform (Bio-Rad) and analyzed using CFX Manager software (v. 3.1; Bio-Rad). Gene expression was quantified using pre-validated PrimePCR primers (Bio-Rad) for *Nox1* (qMmuCED0048182) and *Duox2* (qMmuCID0022771) and the ΔΔCq method with data normalized to the reference genes *Gapdh* (forward primer: 5’-GGAGAAACCTGCCAAGTATG-3’; reverse primer: 5’-GGAGTTGCTGTTGAAGTCG-3’) and *Rplpo* (forward primer: 5’-GCGACCTGGAAGTCCAACTA-3’; reverse primer: 5’-TCTCCAGAGCTGGGTTGTTT-3’).

RNA sequencing was performed by the Genomics and Bioinformatics Core Facility platforms at Sahlgrenska Academy, University of Gothenburg. The quality of isolated RNA was determined using a Bioanalyzer (Agilent) with minimum acceptable RIN value of 8. cDNA was prepared using the TruSeq Stranded Total RNA Sample Preparation kit with Ribo Zero Gold (Rev. E; Illumina) according to manufacturer’s protocol, and sequenced using a NovaSeq 6000 platform (Illumina). Quality of raw sequencing data was assessed using FastQC (v 0.11.2) (http://www.bioinformatics.babraham.ac.uk/projects/fastqc) and if required, Fastq files were quality filtered using prinseq (v 0.20.3) (*45*). Reads were mapped against *Mus musculus* reference genome mm10 using STAR (v 2.5.2b) (*46*). The alignment quality was assessed using samtools (v 1.6) and qualimap (v 2.2.1) (*47, 48*). The number of mapped reads towards annotated features in the reference genome was calculated using HTseq (v 0.5.3p3) (*49*).

### RNA sequencing data analysis

Data analysed in this study included bulk RNA sequencing data generated in the current project supplemented with published bulk RNA sequencing data from sorted colonic goblet cells and enterocytes and single cell RNA sequencing data of sorted colonic goblet cells generated as previously described (*23*) and deposited under GEO accession number GSE144436.

Statistical analysis and identification of differential gene expression between experimental groups based on bulk RNA sequencing data was performed using DESeq2 (*50*) in R. Genes with significant age-correlated monotonic expression patterns were identified by aligning sample age (days *postpartum*) with size factor normalised read counts generated by DESeq2 and calculating Spearman’s rank correlation coefficient. Significant positive and negative age:gene expression correlations were determined after Benjamini-Hochberg FDR correction. For single cell RNA sequencing analysis, demultiplexing, barcoded processing, gene counting and aggregation were performed using Cell Ranger Cell Ranger software (v. 2.1.1) and analysed using Loupe Browser (v 8.0.0; 10X Genomics). Barcodes were filtered to remove non-goblet cells based on zero reads of *Ptprc* (immune cells) or *Chgb* (enterendocrine cells) with *Muc2* reads >3 to avoid enterocyte contamination. Barcodes were further filtered to remove cells with <200 or >5500 UMIs and mitochondrial gene counts >20%, resulting in 6123 total cells post-filtering. Cells were clustered based on K-Means (*K* = 10) and visualised by UMAP embedding. Goblet cell cluster identities were defined based on previous analysis and annotation of the same dataset (*23*). For mapping of *Rattus norvegicus* genes to *Mus musculus* orthologs in single cell RNA sequencing data, *R. norvegicus* gene identifiers were mapped to *M. musculus* identifiers using the g:Orth function of g:Profiler (*51*).

### Microbiota DNA extraction, 16S rRNA gene quantification and sequencing

Faecal and tissue (intestinal and lymph node) DNA was extracted using QIAmp PowerFecal Pro kits (Qiagen) with 4x rounds of 4.5 m/s for 40 s bead-beating using a Fast-Prep System (MPBio). Prior to extraction, tissue cells were lysed by brief processing with an Ultra-Turrax rotor-stator homogeniser, followed by pelleting of bacterial cells and tissue debris by centrifugation at 10,000 RCF for 10 min and discarding the supernatant. For absolute quantification of bacterial 16S rRNA gene copy number, DNA extractions were analyzed by qPCR using SsoFast EvaGreen Supermix (Bio-Rad) with 0.3 µm universal 16S primers 926f (5ʹ-AAACTCAAAKGAATTGACGG-3ʹ) and 1062r (5ʹ-CTCACRRCACGAGCTGAC-3ʹ) with 45 ng template DNA. Reactions were performed and monitored using a CFX96 platform (Bio-Rad). Absolute bacterial 16S copy number was quantified using standard curves generated from qPCR of whole 16S gene amplicons purified from *E. coli*, and data was normalised to initial faecal sample mass. For some experiments, the relative proportion of different major bacterial phyla (*Bacteroidota*, *Firmicutes*, *Proteobacteria*) was determined using taxon specific primers and PCR conditions as previously described (*52*).

For 16S rRNA gene sequencing, extracted faecal DNA was first subjected to PCR amplification, targeting the V5 and V6 hypervariable regions of the 16S rRNA gene using bacteria-specific primers, with sequences previously described: forward 5’-CCATCTCATCCCTGCGTGTCTCCGACTCAGC-barcode-ATTAGATACCCYGGTAGTCC-3’ and reverse 5’-CCTCTCTATGGGCAGTCGGTGATA CGAGCTGACGACARCCATG-3’ (*53*). The thermal cycling conditions were set as follows: an initial denaturation at 94°C for 5 minutes, followed by 35 cycles consisting of denaturation at 94°C for 1 minute, annealing at 46°C for 20 seconds, and elongation at 72°C for 30 seconds, with final elongation phase at 72°C for 7 minutes.

PCR amplicons were resolved using a 1% (w/v) agarose gel, displaying an anticipated amplicon size of approximately 350 base pairs. Following electrophoresis, the amplicons were purified utilizing the QIAQuick Gel Extraction Kit (Qiagen, Dusseldorf, Germany). Quantification of the purified amplicons was performed using the Qubit dsDNA HS Assay Kit on the Qubit 3.0 Fluorometer (ThermoFisher Scientific). For the preparation of template-positive Ion PGM™ Template OT2 400 Ion Sphere™ Particles (ISPs) harbouring clonally amplified DNA, we employed the Ion OneTouch™ Instrument alongside the Ion PGM™ Template OT2 400 Kit (ThermoFisher). Subsequently, sequencing was executed using the Ion PGM™ Sequencing 400 Kit and Ion 316™ Chip V2, within the framework of the Ion PGM™ System (Thermo Fisher) (*54*).

### 16S rRNA gene sequencing data analysis

Fastq sequencing files produced by the Ion Torrent PGM™ System were imported into the Quantitative Insights into Microbial Ecology 2 (QIIME2) version 2018.8.1 framework (https://qiime2.org/) (*55*) and analysed as previously described (*56*). Within this pipeline, Amplicon Sequence Variants (ASVs) were defined at a 97% sequence identity threshold using the standard settings of QIIME2. Taxonomic classification of these ASVs was performed utilizing the q2-feature-classifier plugin paired with a Naïve Bayes classifier. The taxonomic classifications were assigned based on their sequence similarity in the SILVA database (https://www.arb-silva.de/).

The features table and mapping file were used to generate a phyloseq object in R package phyloseq (v 3.4) (*57*). The assessment of alpha diversity (Simpson, and Shannon indices) and beta diversity (Bray-Curtis genus-level community dissimilarities on Principal Coordinates Analysis (PCoA)) were conducted. Statistical analyses of the clustering patterns were performed using the Mann-Whitney U test for alpha diversity measures, and the Adonis test (a type of PERMANOVA) for beta diversity, to ascertain the robustness and statistical significance of the group distinctions using identical distance metrics within phyloseq in R. The LEfSe method was used to compare significant differences in taxa between groups (*58*).

### DSS induced colitis

Colitis was induced in mice by provision of 3% (w/v) dextran sodium sulphate *ad libitum* in drinking water for up to 8 days. At sacrifice, colonic and lymphatic (caudal lymph nodes and spleen) tissues were collected and weighed, and colon length was recorded. The most distal 0.5 cm of colonic tissue was separated and snap frozen for 16S rRNA gene quantification (see above). The remaining colon was flushed, opened longitudinally, and rolled into swiss rolls before fixation in formalin. Fixed swiss rolls were paraffin embedded, sectioned and Alcian blue/Periodic Acid Schiff stained as described above. Histopathological assessment of mid-distal colon tissue swiss roll tissue sections was performed based on an adaptation of a previously defined scoring system assessing inflammatory infiltrate, goblet cell loss, crypt hyperplasia, muscle thickening, submucosal inflammation, abscess formation and ulceration (*59*). Histology scores were generated by averaging independent scores from two blinded assessors, and were summed to generate combined histology scores.

