## Supplemental figures for "Neonatal microbiota colonization drives maturation of primary and secondary goblet cell mediated protection in the pre-weaning colon"

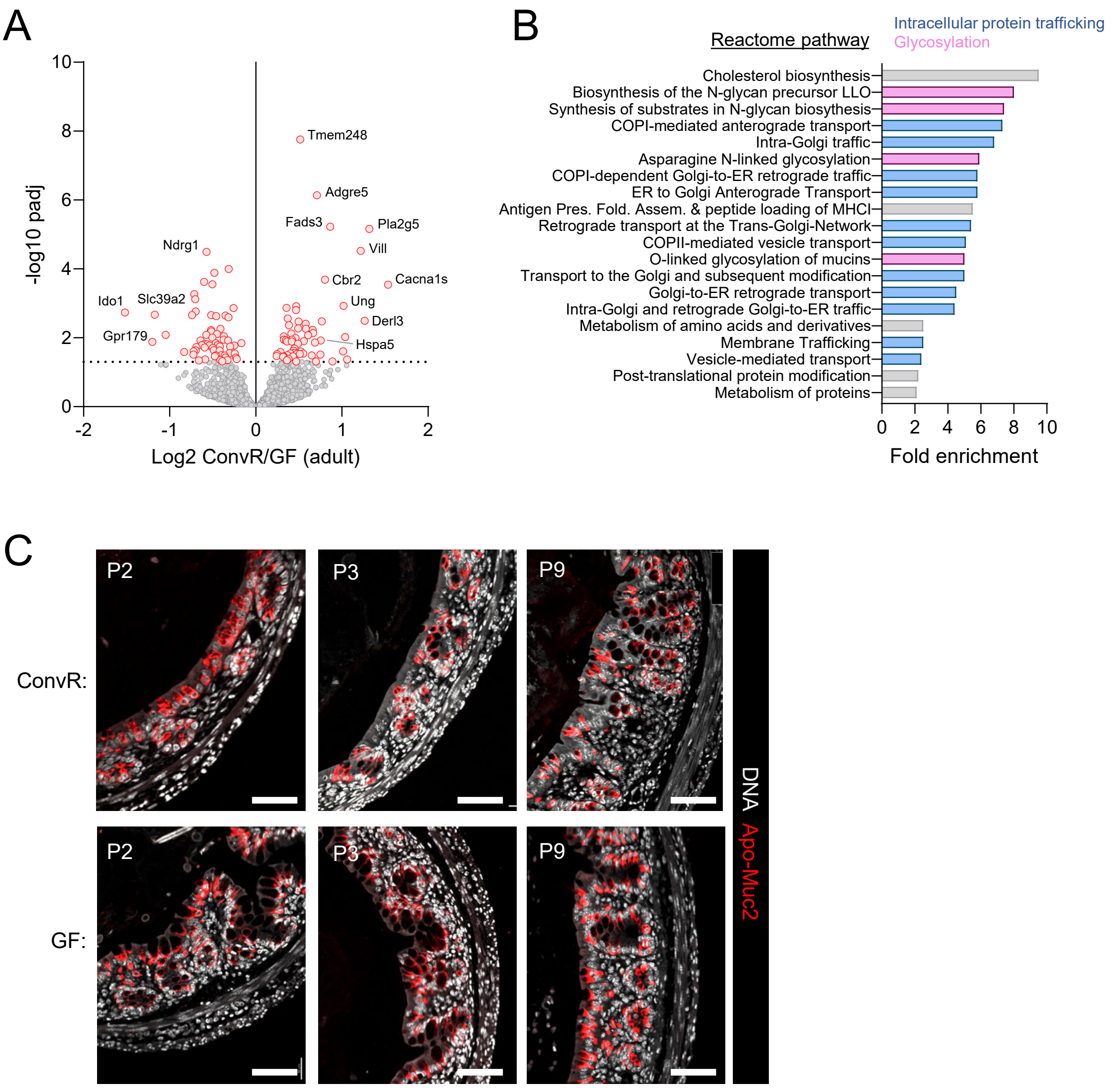


***Supplemental figure 1 (related to Figure 3):***

A) Volcano plot of bulk mRNA sequencing data illustrating GC-enriched genes differentially expressed between ConvR and GF adult colon.

B) Reactome pathways significantly enriched in microbiota-regulated GC genes identified as positively correlated with postnatal age.

C) Confocal micrographs of fixed colonic tissue from adult ConvR and GF mice stained for DNA (grey) and Apo-Muc2 (red).

Data represents n=4 (A, B) or n=4-7 (C) animals per group. All data is pooled from at least 2 independent litters or experiments. Image scale bars are 50µm.

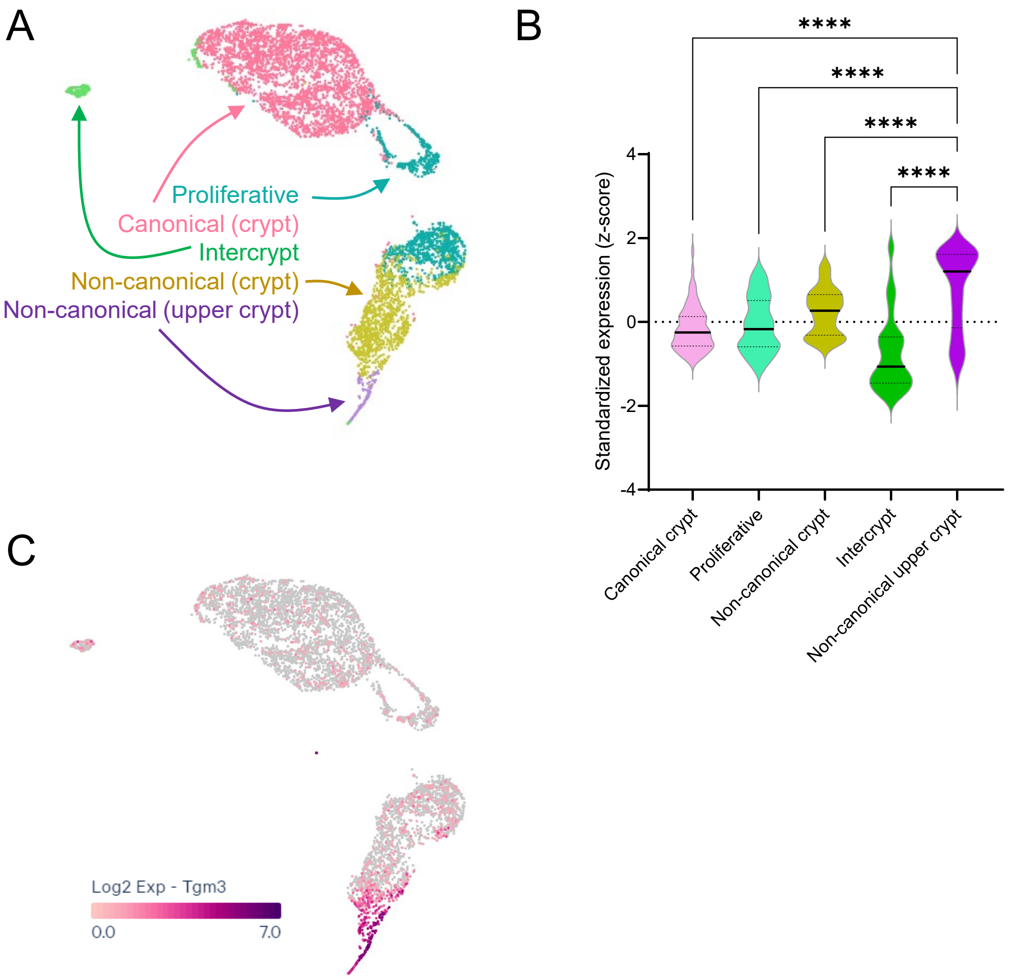


***Supplemental figure 2 (related to Figure 4):***

A) UMAP plot of scRNA-sequencing data from isolated colonic GCs highlighting previously annotated GC clusters (*23*).

B) Per GC cluster standardized gene expression (z-scores) of secreted proteins (n=72) identified as significantly enriched in the P9-Adult compared to P1-P7 colonic mucus proteome (see Fig. 1G, 1H, 4G).

C) UMAP plot illustrating $\log_2$ expression of *Tgm3* in colonic GC scRNA-sequencing data.

Data aggregated from n=2 independent experiments. Violin plot (B) shows median and interquartile range. Statistical comparisons between clusters by one-way ANOVA and Fishers LSD (I); p<0.0001 (****).
